# CerS2 is a druggable target in triple-negative breast cancer

**DOI:** 10.1101/2025.08.15.670525

**Authors:** Hissah Alatawi, Haritha H. Nair, Lingbao Ai, Iqbal Mahmud, Abhishek Gour, Amandeep Singh, Doohyun Baek, Bowen Yan, Chandra K. Maharjan, Weizhou Zhang, Brian K. Law, Maria Zajac-Kaye, Christopher Vulpe, Olga A. Guryanova, Arun K. Sharma, Timothy J. Garrett, Abhisheak Sharma, Coy D. Heldermon, Sukwon Hong, Satya Narayan

**Affiliations:** Department of Anatomy and Cell Biology, College of Medicine, University of Florida, Gainesville, FL 32610; Department of Medicine, College of Medicine, University of Florida, Gainesville, FL 32610; Department of Pathology, Immunology and Laboratory Medicine, College of Medicine, University of Florida, Gainesville, FL 32610; Department of Bioinformatics and Computational Biology, Metabolomics Core Facility, The University of Texas MD Anderson Cancer Center, Houston, TX 77030; Department of Pharmaceutics, College of Medicine, University of Florida, Gainesville, FL 32610; Department of Pharmacology, Penn State Cancer Institute, Hersey, PA 17033; Department of Chemistry, University of Wisconsin-Madison, Madison, Wisconsin 53706; Department of Pharmacology and Therapeutics, College of Medicine, University of Florida, Gainesville, FL 32610; Department of Physiological Sciences, College of Veterinary Medicine, University of Florida, Gainesville, FL 32610; Department of Chemistry, University of Florida, Gainesville, FL 32611; Department of Chemistry, Gwangju Institute of Science and Technology (GIST), 123 Cheomdan-gwagiro, Buk-gu, Gwangju 61005, Republic of Korea

**Keywords:** Triple-negative breast cancer (TNBC), DH20931, ceramide synthase 2 (CerS2), very long chain ceramides (VLCC), ER stress, IP3R1, calcium signaling, mitochondria, apoptosis, targeted therapy

## Abstract

Triple-negative breast cancer (TNBC) remains a significant clinical challenge due to its aggressive nature and lack of effective targeted therapies. The enzyme ceramide synthase 2 (CerS2), which synthesizes pro-apoptotic very long-chain ceramides (VLCCs), represents a promising therapeutic target. Here, we identify and characterize DH20931, a novel, first-in-class small-molecule agonist of CerS2. We demonstrate that DH20931 directly activates CerS2 with nanomolar potency, leading to significant VLCC accumulation in breast cancer cells. This lipotoxic event induces endoplasmic reticulum (ER) stress and triggers apoptosis via the canonical ATF4/CHOP/PUMA signaling pathway. Mechanistically, we uncover a novel interaction between CerS2 and the ER calcium channel, Inositol 1,4,5-trisphosphate receptor 1 (IP3R1). We demonstrate that DH20931 promotes this interaction, enhancing ER-mitochondria proximity and facilitating a CerS2-dependent flux of calcium (Ca²⁺) from the ER into mitochondria. This subsequent mitochondrial Ca²⁺ overload serves as a critical trigger for apoptosis. In preclinical evaluations, DH20931 potently inhibited the growth of TNBC cells in 2D and 3D cultures and significantly suppressed tumor progression in orthotopic and patient-derived xenograft (PDX) models, all while exhibiting a favorable safety profile. Our findings validate CerS2 as a druggable target in TNBC and establish a novel therapeutic strategy that leverages a coordinated attack on cancer cells through ER stress and calcium-mediated mitochondrial dysfunction.

**Graphical Abstract:** 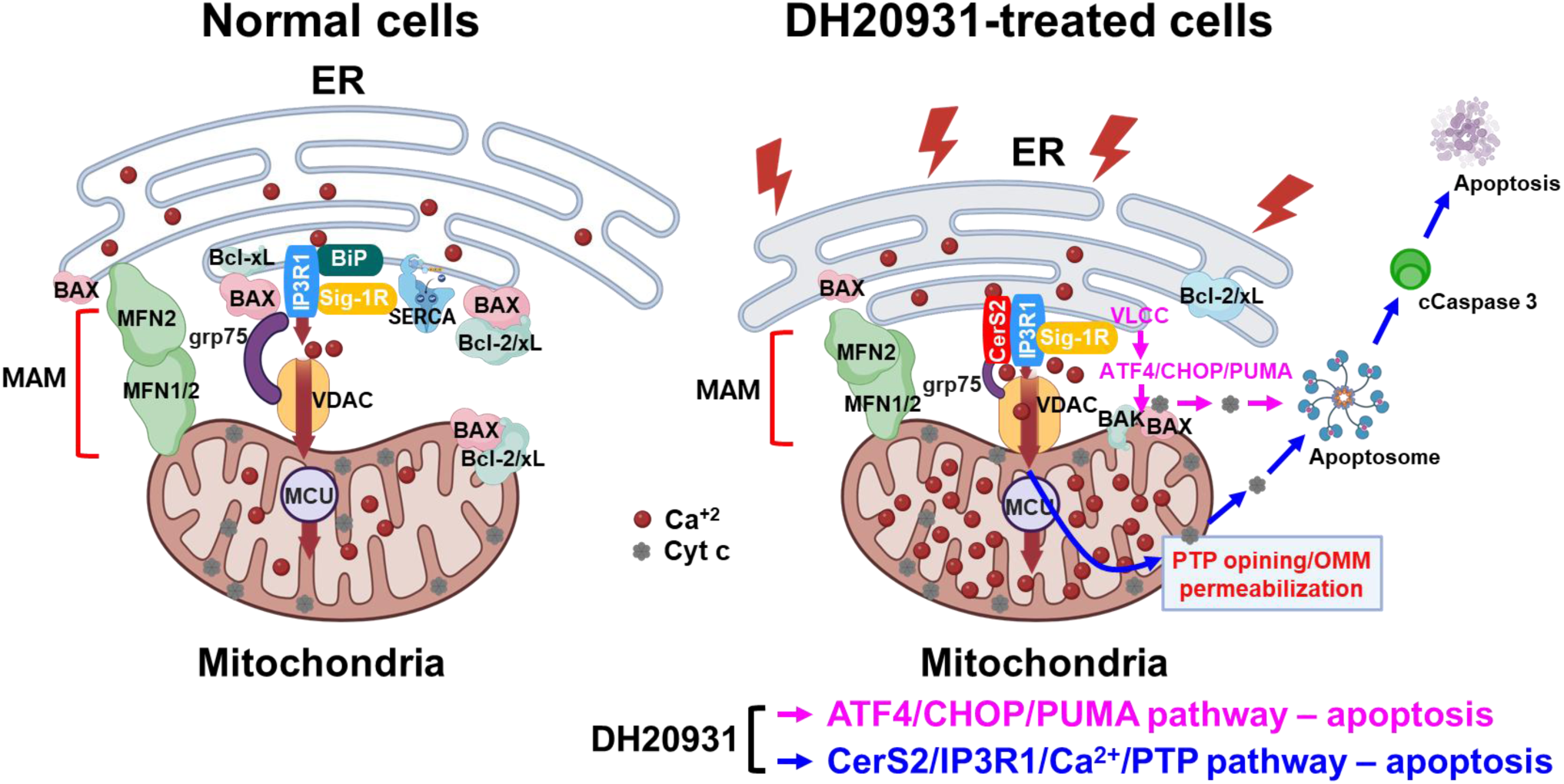

**Highlights:**

- DH20931 is a first-in-class, potent agonist of Ceramide Synthase 2 (CerS2).
- CerS2 activation induces ER stress and engages the ATF4/CHOP/PUMA apoptotic pathway.
- DH20931 promotes a novel CerS2-IP3R1 interaction, causing lethal mitochondrial calcium overload.
- Targeting CerS2 shows significant preclinical efficacy against triple-negative breast cancer.

## INTRODUCTION

Breast cancer is a heterogeneous disease and a leading cause of cancer-related mortality in women worldwide.^1^ The advent of targeted therapies against the estrogen receptor (ER), progesterone receptor (PR), and human epidermal growth factor receptor 2 (HER2) has revolutionized treatment for hormone receptor-positive (HR+) and HER2-positive subtypes. However, a formidable challenge persists in treating triple-negative breast cancer (TNBC), a subtype defined by the lack of ER, PR, and HER2 expression that accounts for approximately 15-20% of all cases.^2^ TNBC is characterized by an aggressive clinical course, including high histological grade, high mitotic indices, a propensity for early relapse peaking within three years of diagnosis, and a distinct pattern of metastasis to visceral organs such as the brain and lungs.^3^ Due to the absence of well-defined molecular targets, systemic chemotherapy remains the primary treatment modality.^3^ While newer agents—including immune checkpoint inhibitors for tumors expressing PD-L1, PARP inhibitors for patients with germline *BRCA1/2* mutations, and antibody-drug conjugates (*e.g.,* Trastuzumab Emtansine, Trastuzumab Deruxtecan, Sacituzumab Govitecan)—have demonstrated benefit in specific subsets, a large proportion of patients do not respond or rapidly develop resistance.^2–7^ This therapeutic impasse, compounded by the significant molecular heterogeneity within TNBC itself, underscores a critical and urgent unmet clinical need for novel therapeutic targets and strategies that can combat this recalcitrant disease independent of classical receptor status.

Cellular metabolism is increasingly recognized as a hallmark of cancer, with malignant cells undergoing metabolic reprogramming to support their uncontrolled proliferation and survival.^8^ Among the most altered pathways are those involved in lipid metabolism. Sphingolipids, a complex class of lipids, have emerged from their historical role as simple structural components of cell membranes to be appreciated as critical signaling molecules that regulate a diverse array of cellular processes, including proliferation, apoptosis, and senescence.^9^ The balance between pro-apoptotic ceramides and pro-survival sphingosine-1-phosphate (S1P) acts as a “sphingolipid rheostat,” where a shift towards ceramide accumulation promotes cell death, establishing ceramide-elevating strategies as a valid anti-cancer approach.^9^ The central molecule in this axis, ceramide, is synthesized *de novo* in the endoplasmic reticulum (ER) by a family of six ceramide synthase (CerS) enzymes, CerS1-6.^10^ Each CerS isoform exhibits remarkable specificity for fatty acyl-CoAs of different chain lengths, thereby producing ceramides with distinct N-acyl chains and, consequently, distinct biological functions.^11^ For instance, CerS1 preferentially synthesizes C18-ceramide, while CerS2 is primarily responsible for the synthesis of very long-chain ceramides (VLCCs) containing C22-, C24-, and C26-acyl chains.^12^

The role of CerS2 in cancer biology has been considered complex and seemingly paradoxical. A substantial body of evidence positions CerS2 as a tumor suppressor.^13,14^ In multiple cancer types, including breast, bladder, and liver cancer, decreased expression of CerS2 is correlated with poor prognosis, increased tumor invasion, and reduced patient survival.^15^ Mechanistically, the VLCCs produced by full-length CerS2 have been shown to suppress cancer cell invasion, in part by inhibiting matrix metalloproteinases.^13^ However, this narrative sometimes is complicated by reports that, under certain conditions,CerS2 mRNA and protein levels are elevated in aggressive tumors promoting proliferation.^16^ This apparent contradiction appears to stem from complex regulatory mechanisms, including alternative splicing. A splice variant of *CERS2* that lacks exon 8—a critical part of the catalytic domain—has been identified in Luminal B breast cancer and is associated with poor prognosis and increased cell migration, as it fails to produce tumor-suppressive VLCCs.^14^ Furthermore, the activity of canonical CerS2 can be directly suppressed by inhibitory binding partners, such as the anti-apoptotic protein Bcl2L13.^17^ This clarifies the therapeutic objective: it is not merely to increase *CERS2* expression, but to specifically enhance the enzymatic activity of the full-length, canonical CerS2 protein to drive VLCC production.

CerS2 and its VLCC products are intrinsically linked to the function and integrity of the ER, the primary site of their synthesis.^18^ Perturbations that disrupt ER homeostasis, such as the accumulation of unfolded proteins or lipid bilayer stress, activate a sophisticated signaling network called the unfolded protein response (UPR).^19^ While initially adaptive, prolonged or irremediable ER stress, mediated by the sensors PERK, IRE1α, and ATF6, switches the UPR into a pro-apoptotic program.^20^ The ATF4 can also function entirely independently of two of the three main ER stress sensors, IRE1 and ATF6 and induce apoptosis through PERK-eIF2α-ATF4-CHOP integrated stress response (ISR). ^21^ The accumulation of VLCCs, by altering the biophysical properties of the ER membrane, is a potent inducer of lipotoxic ER stress, leading to robust activation of the pro-death PERK-eIF2α-ATF4-CHOP pathway.^22–24^ Beyond protein folding, the ER is the cell’s main intracellular calcium (Ca^2+^) reservoir, and its communication with mitochondria is a critical nexus for apoptosis regulation.^25^ This communication occurs at specialized contact sites known as mitochondria-associated membranes (MAMs), which physically tether the two organelles and serve as hubs for both lipid synthesis and Ca^2+^ signaling.^26^ At MAMs, Ca^2+^ released from the ER through channels like the inositol 1,4,5-trisphosphate receptor 1 (IP3R1) is efficiently taken up by mitochondria.^27,28^ While modest mitochondrial Ca^2+^ uptake supports bioenergetics, an excessive and sustained flux triggers the opening of the mitochondrial permeability transition pore (mPTP), leading to the collapse of the mitochondrial membrane potential, release of cytochrome c, and activation of the intrinsic apoptotic cascade.^29,30^

In this study, we identify and characterize DH20931, a novel biisoquinoline derivative, as the first-in-class small-molecule activator of CerS2. We demonstrate that pharmacological activation of CerS2 by DH20931 represents a powerful therapeutic strategy that is effective across all breast cancer subtypes, including hard-to-treat TNBC. We elucidate a dual mechanism of action whereby DH20931-driven VLCC accumulation first initiates lipotoxic ER stress, engaging the pro-apoptotic ATF4/CHOP/PUMA pathway. Concurrently, we reveal a previously unknown mechanism wherein DH20931 promotes direct physical interaction between CerS2 and the Ca^2+^ channel IP3R1 at MAMs. This novel interaction facilitates a lethal flux of Ca^2+^ from the ER to the mitochondria, triggering catastrophic mitochondrial-dependent apoptosis. These findings not only validate CerS2 as a bona fide druggable target in breast cancer but also uncover a new layer of crosstalk between sphingolipid metabolism, ER stress, and calcium signaling in the control of cell death.

## RESULTS

### Discovery of DH20931, a novel activator of ceramide synthase 2

To identify novel therapeutic strategies for breast cancer based on metabolic vulnerabilities, we investigated the effects of DH20931, a proprietary small molecule, on the cancer cell lipidome. Using untargeted lipidomics, we analyzed extracts from the MDA-MB-468 TNBC cell line and BT-474 (Luminal B) tumor xenografts treated with DH20931. The analysis revealed a distinct lipid class alteration in the ceramide profile of treated samples compared to vehicle-treated controls (Figure 1A). The most prominent change was an increase in ceramide species containing very long-chain fatty acids (VLCCs), specifically those with C22:0, C24:0, C24:1, and C26:0 N-acyl chains. This specific enrichment implicated ceramide synthase 2 (CerS2), the enzyme responsible for synthesizing this class of ceramides^12^, as a potential molecular target of DH20931.

**Figure 1.**
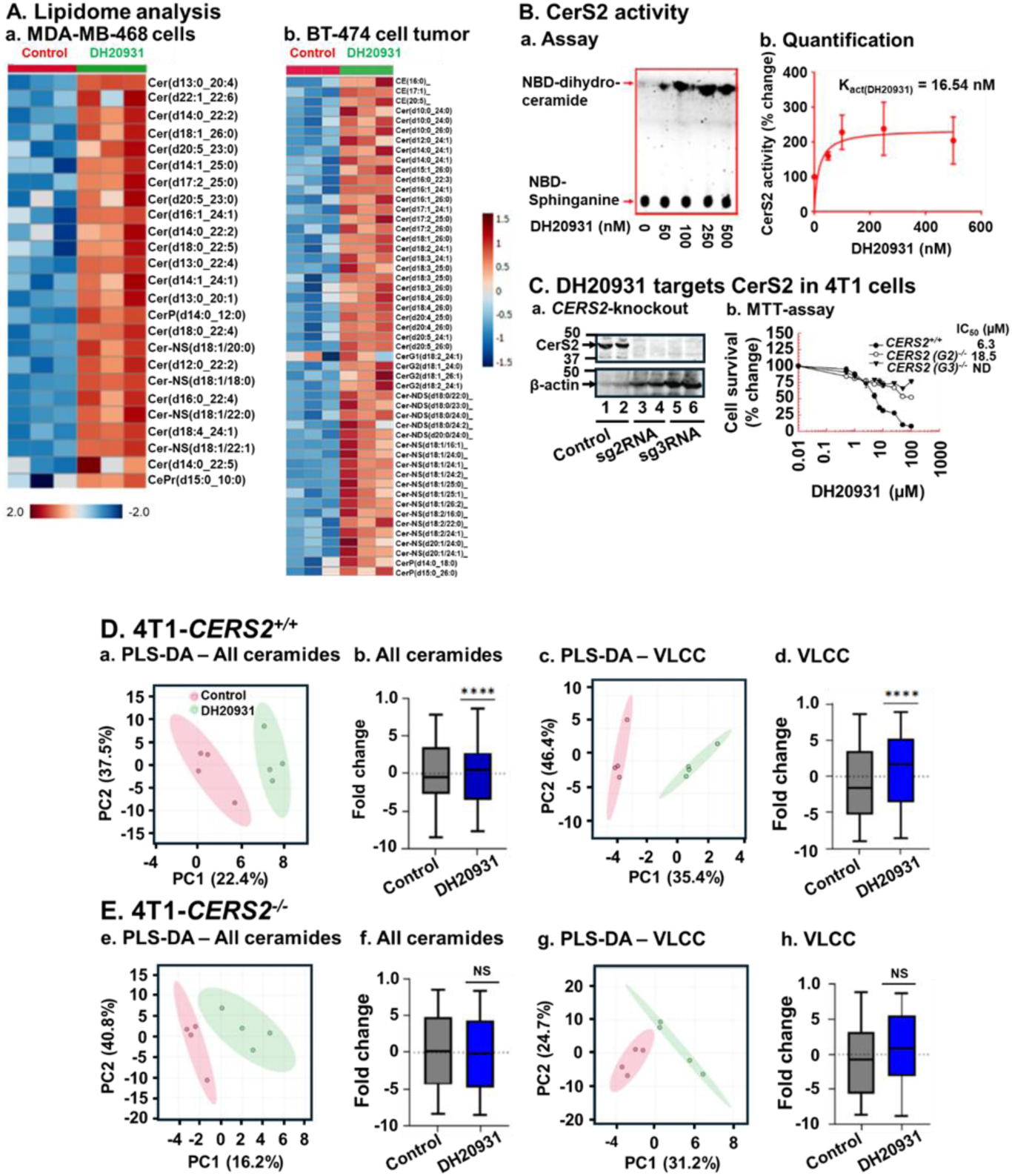
DH20931 Enhances Ceramide Synthase 2 (CerS2) Activity and Very Long-Chain Ceramide (VLCC) Synthesis in Breast Cancer Cells. **(A)** Untargeted lipidome analysis of MDA-MB-468 and BT-474 tumor xenografts from control and DH20931-treated female NDG mice. Data are representative of three biological replicates per group. **(B)** Direct induction of CerS2 enzymatic activity by DH20931. (a) A representative thin-layer chromatography (TLC) gel showing the formation of NBD-C6-dihydroceramide. (b) Quantification of NBD-C6-dihydroceramide levels from three independent experiments. Data are presented as mean ± SEM. **(C)** The biological activity of DH20931 is dependent on CerS2. (a) Western blot analysis confirming CRISPR-Cas9-mediated knockdown of CerS2 in 4T1 cells. (b) Cell viability curves for 4T1 cells treated with DH20931, demonstrating that CerS2 is required for drug-mediated sensitization. The IC₅₀ values were determined by MTT assay. Data are representative of four independent experiments. **(D)** Lipidome analysis of 4T1-*CERS2*⁺^/^⁺ and 4T1-*CERS2^−/-^*cells. Wild-type (4T1-*CERS2*⁺^/^⁺) and CerS2-knockout (4T1-*CERS2*⁻^/^⁻) cells were treated for 24 h with vehicle control or 10 µM DH20931. A total of 81 ceramide species, including 46 VLCC species, were quantified. (a–d) Partial Least Squares-Discriminant Analysis (PLS-D) and box-and-whisker plots of total ceramide and VLCC levels in 4T1-*CERS2*⁺^/^⁺ cells. (e– h) PCA and box-and-whisker plots of total ceramide and VLCC levels in 4T1-*CERS2*⁻^/^⁻ cells. Data are presented as mean ± SEM from four independent samples. NS, not significant; ****p < 0.0001. See also Figure S2.

To test this hypothesis directly, we performed an *in vitro* enzymatic assay using cell lysates from cells overexpressing CerS2 in HEK293T cells.^31^ The assay measures the incorporation of a fluorescent NBD-sphinganine substrate with a fatty acyl-CoA to form NBD-ceramide. Treatment with DH20931 resulted in a dose-dependent increase in CerS2 activity, as visualized by the increased formation of fluorescent product on TLC plates (Figure 1B, panel a). Quantification of this activity revealed that DH20931 is a remarkably potent activator of CerS2, with a calculated activation constant (K_act(DH20931)_) of 16.54 nM (Figure 1B, panel b). To our knowledge, DH20931 is the first small-molecule direct agonist of CerS2 to be described, representing a novel chemical probe and potential therapeutic agent.

To determine whether *CERS2* expression level is linked with the prognosis of breast cancer, we performed Kaplan-Meier survival analysis.^32^ Our results clearly showed a longer survival for breast cancer patients with higher levels of *CERS2* expression (Figure S1). While this correlation was statistically significant across multiple breast cancer subtypes, including All-BRCA (all breast cancer), Luminal A, Luminal B, ER+, and HER2+, it was not significant in Basal-like (TNBC) breast cancer (Figure S1). This suggests that the prognostic value of *CERS2* expression is highly dependent on the specific molecular characteristics of the tumor. In TNBC, which is characterized by a more aggressive phenotype and a lack of targeted receptors, other pro-survival pathways may override the tumor-suppressive effects of *CERS2*. In this study, we have shown that DH20931 can significantly induce the sensitization of all breast cancer subtypes, including TNBC. This finding is particularly important because it suggests that while baseline *CERS2* expression may not be a reliable prognostic marker for TNBC, actively stimulating the CerS2-mediated ceramide pathway with a compound like DH20931 can still provide therapeutic benefits. This demonstrates a potential strategy to overcome the inherent resistance of TNBC by directly targeting a pathway that promotes cell death, even when the pathway’s basal activity is not correlated with survival outcomes.

To establish that CerS2 is the critical mediator of DH20931’s biological effects in cancer cells, we employed a genetic approach. Using CRISPR/Cas9, we generated a stable knockout of the *CERS2* gene in the highly aggressive 4T1 mouse TNBC cell line (4T1-*CERS2*⁻^/^⁻), confirming the absence of the protein by Western blot (Figure 1C, panel a). We then compared the sensitivity of these cells to their wild-type counterparts (4T1-*CERS2*⁺^/^⁺) in cell survival assays. As shown in Figure 1C, panel b, while DH20931 potently inhibited the viability of wild-type 4T1 cells, the *CERS2*-knockout cells were almost completely resistant to the drug, exhibiting a greater than 10-fold increase in the IC₅₀ value. This desensitization was further confirmed in a long-term clonogenic survival assay, where DH20931 effectively reduced the colony-forming ability of wild-type cells but had minimal impact on the *CERS2*⁻^/^⁻ cells (Figure S2).

Finally, to link the biological effect directly to the compound’s enzymatic action, we performed lipidomic analysis on the wild-type and knockout 4T1 cells. As expected, treatment of 4T1-*CERS2*⁺^/^⁺ cells with DH20931 for 24 h led to an accumulation of total ceramides, which was driven by a significant increase in VLCCs (Figure 1D, panels b-d). Partial Least Squares-Discriminant Analysis (PLS-DA) of the lipidome showed a clear separation between vehicle- and DH20931-treated wild-type cells, further confirmed by PLS-DA performance value >80% (Figure 1D, panel a). In stark contrast, DH20931 treatment had no or minimal effect on the ceramide or VLCC content of 4T1-*CERS2*⁻^/^⁻ cells, further verified by PLS-DA performance of <70% (Figure 1E, panels f-h), and PLS-DA analysis showed less profile distinction between the treatment groups (Figure 1E, panel e). Taking together, these data provide genetic and biochemical evidence that DH20931 exerts its cytotoxic effects by directly activating CerS2 and driving the overproduction of pro-apoptotic VLCCs.

### Computational modeling predicts a direct and stable interaction between DH20931 and CerS2

To gain insight into the molecular basis of DH20931’s potent activation of CerS2, we turned to computational modeling. Since no experimental structure of CerS2 currently exists, we utilized the high-quality predicted structure of human CerS2 from the AlphaFold database (AF-Q3ZBF8-F1-model_v4) as our receptor model. We first identified potential binding pockets on the protein surface and then performed Induced Fit Docking (IFD) with DH20931. This advanced docking protocol allows for flexibility in both the ligand and the protein’s active site residues, providing a more realistic model of the binding event. The resulting poses were then re-scored using MM-GBSA (Molecular Mechanics Generalized Born Surface Area) calculations to estimate the binding free energy. The top-ranked pose demonstrated an exceptionally strong predicted binding free energy of –92.86 kcal/mol, suggesting a highly stable and high-affinity interaction (Figure 2A, panel a).

**Figure 2.**
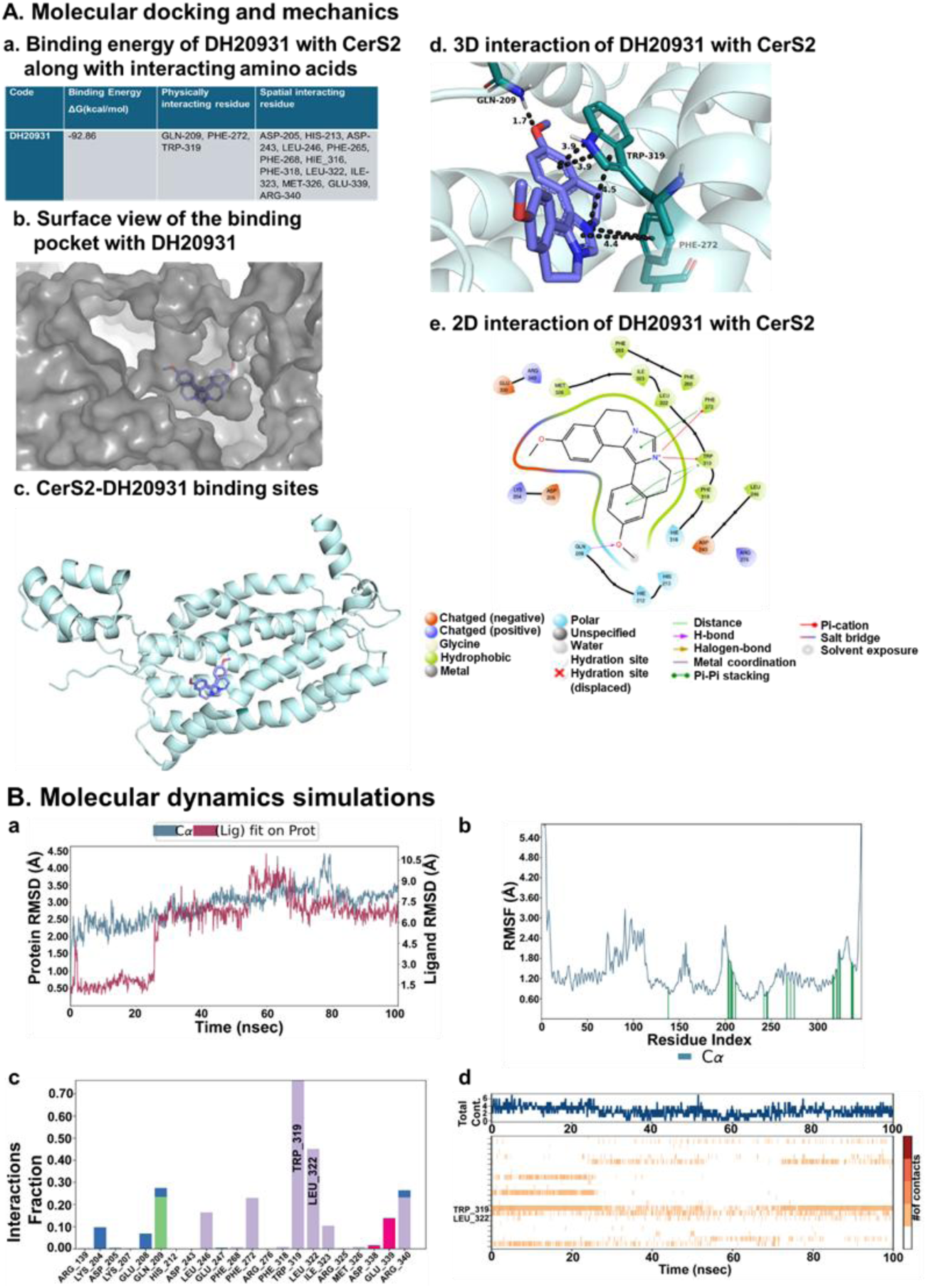
Molecular Modeling of the DH20931 and CerS2 Interaction. **(A)** Induced-fit docking model of DH20931 with the CerS2 protein. (a) A summary of the predicted binding energy and key interacting amino acid residues. (b) A surface-view representation of the CerS2 binding pocket occupied by DH20931. (c) A ribbon model of the full-length CerS2 protein highlights the DH20931 binding site. (d and e) 3D and 2D representations illustrating the specific interactions between DH20931 and the binding pocket of CerS2. **(B)** Molecular dynamics (MD) simulations of the DH20931-CerS2 complex over a 100 ns trajectory. (a) Root-mean-square deviation (RMSD) of the complex over time. (b) Root-mean-square fluctuation (RMSF) of individual amino acid residues. Green lines indicate residues in contact with DH20931. (c) Normalized stacked bar chart quantifying the types of protein-ligand interactions (hydrogen bonds, ionic, water bridges, and hydrophobic) throughout the simulation. A value of 1.0 indicates an interaction is maintained for 100% of the trajectory. (d) A timeline detailing the specific protein-ligand contacts over the course of the simulation.

Detailed analysis of this binding pose revealed a snug fit of DH20931 within a deep, hydrophobic pocket of the enzyme (Figure 2A, panel b). The stability of the complex was mediated by a network of specific non-covalent interactions (Figure 2A, panels c-e). Notably, the binding was anchored by a π–cation interaction between the positively charged quaternary nitrogen of DH20931’s imidazole ring and the aromatic ring of residue PHE-272 and five membered-ring of residue TRP-319. This was further stabilized by extensive π–π stacking interactions between the aromatic systems of DH20931 and the indole ring of TRP-319, as well as additional contacts with PHE-272. Additionally, methoxy group of the ligand made a hydrogen bond with residue GLN-209 with a bond length of 1.7 Å to further stabilize the complex.

To assess the stability of this predicted binding mode over time, we performed a 100-nanosecond all-atom molecular dynamics (MD) simulation of the DH20931-CerS2 complex. The root-mean-square deviation (RMSD) of the protein’s Cα atoms remained stable within the accepted range of 2.3–3.5 Å throughout the simulation, indicating that the binding of DH20931 did not induce major conformational instability in the protein’s overall fold (Figure 2B, panel a, blue line). The ligand itself, after an initial equilibration period, also settled into a stable conformation within the binding pocket, with its RMSD stabilizing for the final 35 ns of the simulation (Figure 2B, panel a, red line). Furthermore, analysis of the root-mean-square fluctuation (RMSF) of individual residues showed that the residues lining the binding pocket exhibited low fluctuation, indicating they were stably engaged in interactions with the ligand (Figure 2B, panel b). Analysis of the protein-ligand contacts throughout the simulation revealed that the interactions observed in the initial docking pose were largely maintained. Hydrophobic contacts, particularly with TRP-319, were the predominant interaction type, with high interaction fractions of 0.80 (Figure 2B, panels c, d). Importantly, the MD simulation also revealed the formation of new, stable interactions not predicted by the static docking model, including a hydrophobic interaction with LEU-322 and a salt bridge with GLU-339, which likely contribute further to the high-affinity binding. Collectively, these computational results provide a strong, atomistic-level rationale for the potent and specific activation of CerS2 by DH20931, predicting a direct, stable, and high-affinity binding event within a well-defined pocket of the enzyme.

### DH20931 inhibits breast cancer growth *in vitro* and *in vivo*, independent of hormone receptor status

Having established CerS2 as the direct target of DH20931, we next evaluated its therapeutic potential across a broad spectrum of breast cancer subtypes. We performed cell viability (MTT) assays on a large panel of human breast cancer cell lines, including Luminal A (T47D), Luminal B (BT-474), HER2-positive (SKBr3), and multiple TNBC models (MDA-MB-468, MDA-MB-231, MDA-MB-436, 4T1). DH20931 demonstrated potent cytotoxicity against all tested cancer cell lines, with IC₅₀ values in the low-micromolar range, irrespective of their ER, PR, or HER2 status (Figure 3A). This broad efficacy underscores a key potential advantage of a CerS2-targeted approach: the ability to bypass the classical drivers of breast cancer that define current targeted therapy paradigms. Crucially, DH20931 showed minimal toxicity toward non-malignant human mammary epithelial cells (HMEpC), indicating a favorable therapeutic window. Its efficacy also extended to patient-derived xenograft (PDX) tissue cell lines, including the HCI-001 (TNBC) and HCI-012 (HER2+) models.^33^

**Figure 3.**
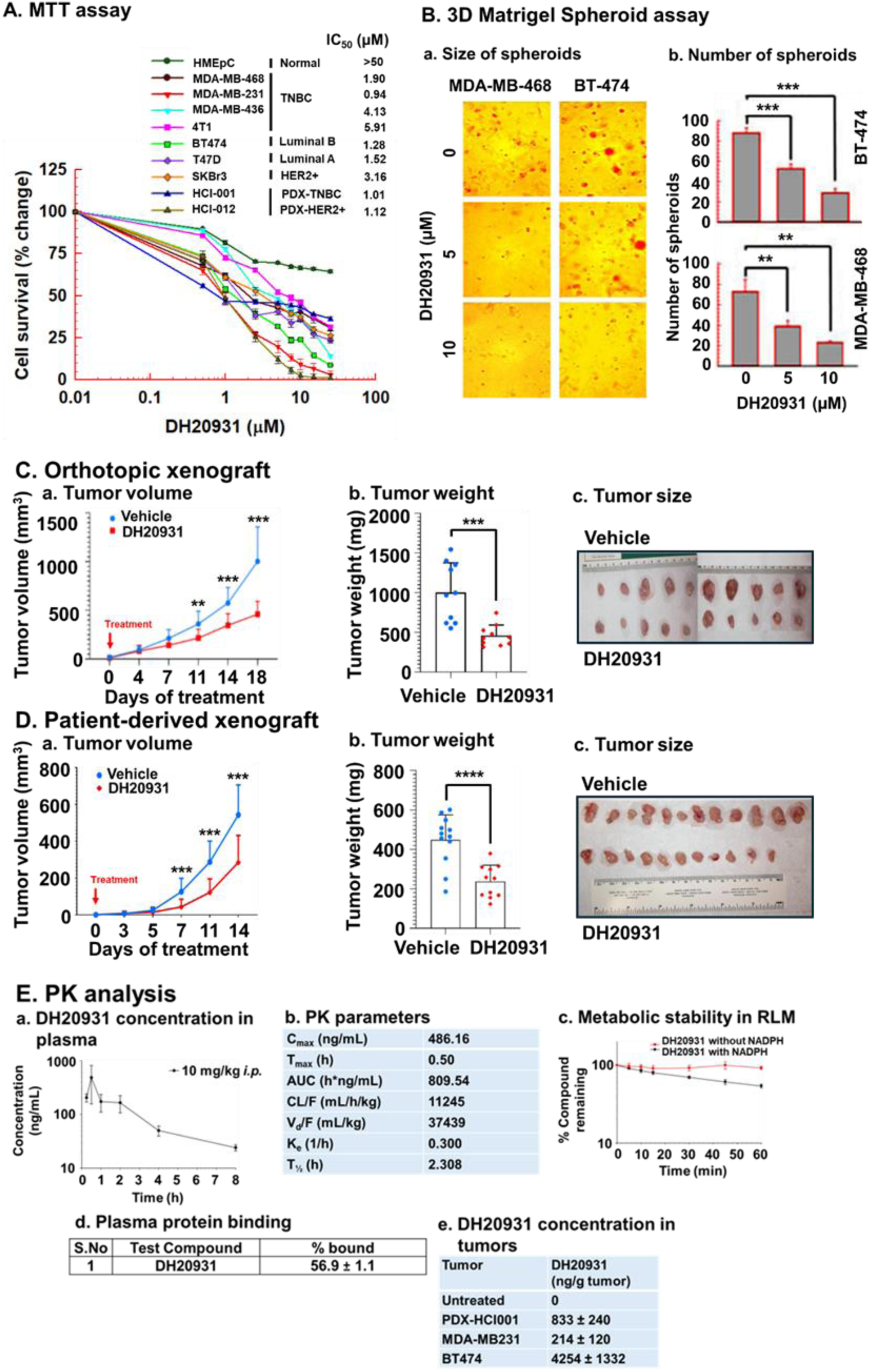
DH20931 Exhibits Antitumor Activity in Breast Cancer Models and a Favorable Pharmacokinetic Profile. **(A)** Cell viability of Luminal A, Luminal B, HER2-positive, and triple-negative breast cancer (TNBC) cell lines after a 72-hour treatment with DH20931, as determined by MTT assay. Data are presented as mean ± SEM from four independent experiments. **(B)** DH20931 inhibits the growth of MDA-MB-468 and BT-474 cells in a 3D Matrigel spheroid assay. Spheroids were treated with 5 or 10 µM DH20931 for 10 days. (a) Representative images of spheroids (10x magnification). (b) Quantification of spheroid number. Data are presented as mean ± SEM. ***p < 0.001. **(C)** DH20931 suppresses tumor growth in an orthotopic MDA-MB-231 TNBC mouse model. Mice (n=10/group) were treated with vehicle or DH20931 (6 mg/kg, 5 days/week). Graphs show (a) tumor volume over time, (b) final tumor weight, and (c) representative images of excised tumors. Data are presented as mean ± SD. **p < 0.01, ***p < 0.001. **(D)** DH20931 inhibits tumor growth in a TNBC patient-derived xenograft (PDX) model (HCI-001). The treatment schedule was identical to (C). Data (n=10-11 mice/group) show tumor volume, final tumor weight, and representative tumor images. Data are presented as mean ± SD. ***p < 0.001, ****p < 0.0001. **(E)** Pharmacokinetic (PK) and biodistribution analysis of DH20931. (a) Plasma concentration-time profile of DH20931 in C57BL/6J mice after a single 10 mg/kg intraperitoneal administration. (b) PK parameters derived from the plasma concentration-time data. (c) Metabolic stability of DH20931 in a rat liver microsome assay. (d) Protein binding of DH20931 in human plasma. (e) Concentrations of DH20931 in tumor tissues from the indicated models. All data are presented as mean ± SD from 3-4 samples per group. See also Figures S3, S4, S5 and Tables S1-S4.

To better mimic the three-dimensional architecture and drug penetration challenges of solid tumors, we tested DH20931 in 3D Matrigel spheroid cultures. When MDA-MB-468 (TNBC) and BT-474 (Luminal B) cells were grown as spheroids, treatment with DH20931 caused a dose-dependent reduction in both the size and number of established spheroids (Figure 3B). This result demonstrates that DH20931 can effectively penetrate multi-layered cell structures and induce cell death, a critical attribute for *in vivo* efficacy.

Encouraged by these *in vitro* findings, we proceeded to evaluate the antitumor activity of DH20931 *in vivo*. We first used an orthotopic xenograft model, implanting MDA-MB-231 TNBC cells into the mammary fat pads of immunodeficient NSG (NOD.Cg-Prkdc^scid^ Il2rg^tm1Wjl^/SzJ) mice. Once tumors were established, mice were treated with either vehicle or DH20931 (6 mg/kg, *i.p.*, 5 days/week). This regimen resulted in statistically significant inhibition of tumor growth compared to the vehicle-treated control group (Figure 3C). At the end of the study, the final weight of excised tumors from the DH20931-treated group was significantly lower than that of the control group. Importantly, this potent efficacy was not limited to a single TNBC model; similar tumor growth inhibition was observed in orthotopic models using MDA-MB-468 (TNBC) and BT-474 (Luminal B) cells (Figure S3), further reinforcing the broad, receptor-independent activity of DH20931.

To assess efficacy in a more clinically relevant setting, we utilized the HCI-001 TNBC patient-derived xenograft (PDX) model. PDX models are derived directly from patient tumors and are known to better recapitulate the heterogeneity and therapeutic response of human cancer.^34^ Following implantation and establishment of HCI-001 tumors in NSG mice, treatment with DH20931 using the same dosing schedule resulted in statistically significant suppression of tumor growth (Figure 3D). This robust efficacy in a PDX model provides strong evidence for the potential clinical translatability of DH20931. Throughout all *in vivo* studies, the therapeutic dose of 6 mg/kg was well-tolerated. We observed no signs of overt toxicity, adverse behavioral changes, or significant loss of body weight (Figure S4). Histopathological analysis of major organs and analysis of blood parameters at the end of the studies revealed no significant organ or major hematological toxicities (Figure S5, Table S1). The observed changes in Mean Corpuscular Volume (MCV), Mean Corpuscular Hemoglobin (MCH), and Mean Corpuscular Hemoglobin Concentration (MCHC) in DH20931-treated mice were subtle and did not lead to a manifest anemia, as evidenced by the consistent red blood cells (RBC), hemoglobin (HGB), and hematocrit (HCT) values. The minor decreases in MCV and MCH, and slight fluctuations in MCHC, suggest a subtle impact on red blood cell maturation or hemoglobinization rather than outright suppression of erythropoiesis or significant red cell destruction (Table S1).

Finally, we characterized the pharmacokinetic (PK) properties of DH20931. Following a single 10 mg/kg intraperitoneal injection in mice, DH20931 exhibited favorable plasma exposure, with a peak concentration (C_max_) of 486.2 ng/mL and an elimination half-life (t_1/2_) of 2.3 h (Figure 3E, panels a, b). The compound also demonstrated good metabolic stability in rat liver microsome assays and moderate binding to human plasma proteins (Figure 3E, panels c, d). The Parallel Artificial Membrane Permeability (PAMPA) assay classified DH20931 in low permeability class of compounds (Table S3). While *in vitro* permeability assay across Caco-2 cell monolayer suggested DH20931 may be a substrate for P-glycoprotein (P-gp) efflux transporters, it was not a substrate for breast cancer resistance protein (BCRP), a transporter often overexpressed in tumors (Table S4). Most importantly, analysis of tumor tissues from our xenograft models confirmed that DH20931 achieves and maintains significant concentrations within the tumor mass, demonstrating excellent biodistribution to its site of action (Figure 3E, panel e).

These experimental findings were supported by *in silico* analysis. The Swiss-ADME tool predicted a molecular weight and topological polar surface area (TPSA, 27.27 Å²) consistent with good membrane permeability. Furthermore, the compound’s LogS and LogP values suggested favorable aqueous solubility and lipophilicity, respectively. Importantly, DH20931 showed no violations of Lipinski’s Rule of Five and triggered no PAINS (Pan-Assay Interference Compounds) alerts, reinforcing its drug-like potential (Table S2).

In summary, DH20931 exhibits potent, broad-spectrum, and receptor-independent antitumor activity *in vitro* and *in vivo*, which is supported by a favorable safety and pharmacokinetic profile, strongly positioning it for further preclinical and clinical development.

### DH20931 induces ER stress through the ATF4 pathway in a CerS2-dependent manner

The accumulation of VLCCs in the ER membrane is a potent form of lipotoxic stress known to disrupt ER homeostasis and trigger the Unfolded Protein Response (UPR). To investigate this, we used Transmission Electron Microscopy (TEM) to visualize the ultrastructural consequences of DH20931 treatment. In both MDA-MB-468 and 4T1-*CERS2*⁺^/^⁺ cells, a 24-hour treatment with DH20931 induced dilation and vesiculation of the ER lumen (Figure 4A, panels a, b, red arrows). This morphology is the classic ultrastructural hallmark of severe ER stress and was comparable to the effects of thapsigargin (Tg), an established pharmacological inducer of ER stress.^35,36^ Critically, these morphological changes were completely absent in 4T1-*CERS2^−/-^* cells treated with DH20931 (Figure 4A, panel c), confirming that the induction of ER stress is a direct consequence of CerS2 activation and VLCC production.

**Figure 4.**
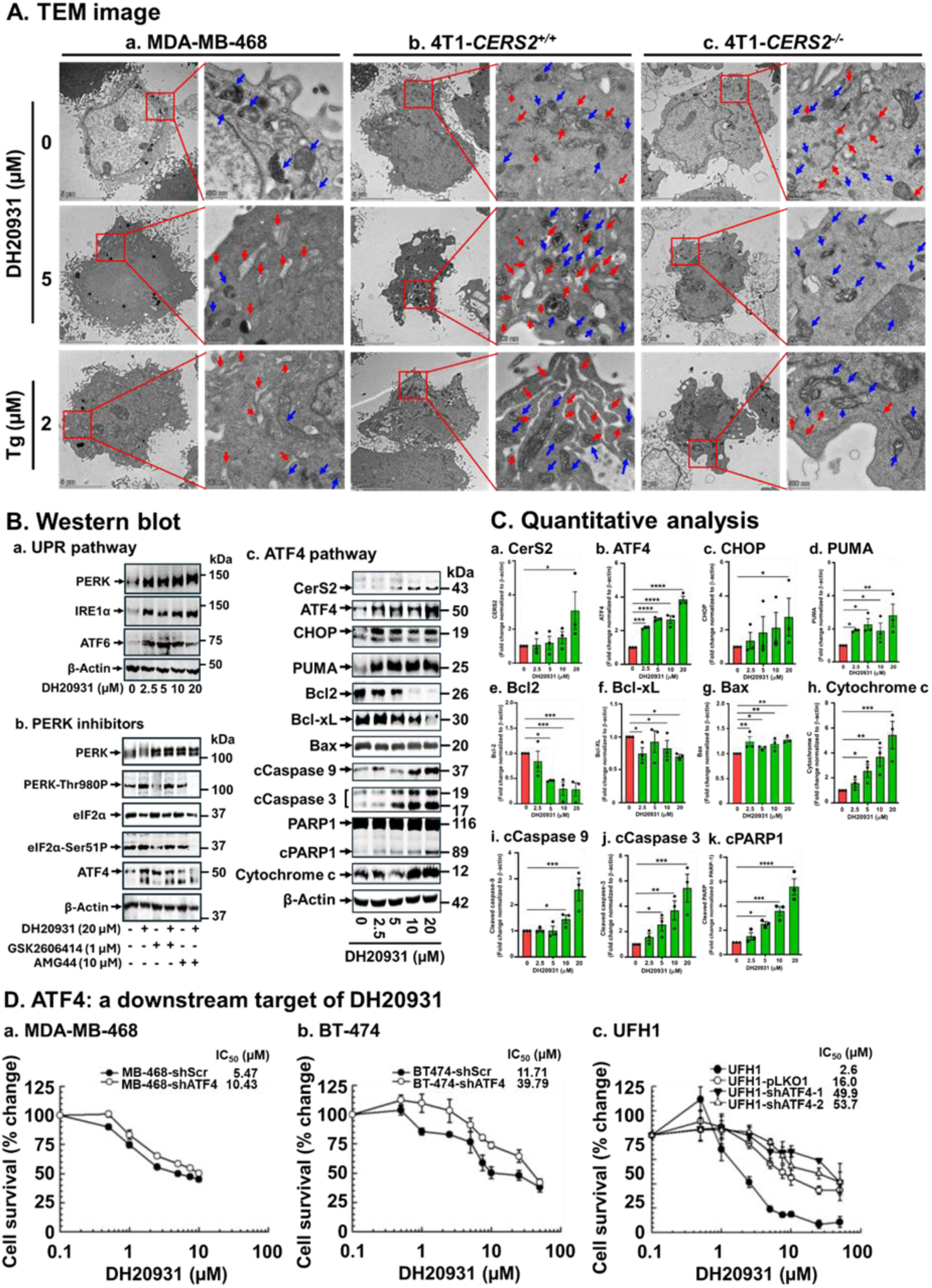
DH20931 Induces ER Stress in 4T1 Cells in a CerS2-Dependent Manner. **(A)** Transmission Electron Microscopy (TEM) images of MDA-MB-468 (a), 4T1-*CERS2*⁺^/^⁺ (b), and 4T1-*CERS2*⁻^/^⁻ (c) cells treated with vehicle, 5 µM DH20931, or 2 µM Thapsigargin (Tg) for 24 h. Blue arrows indicate mitochondria, and red arrows indicate dilated endoplasmic reticulum (ER). Each image is representative of at least two different fields from 4-8 sections. **(B)** Western blot analysis of (a) key UPR proteins and (b) the effect of PERK inhibitors on DH20931-induced phosphorylation of PERK and its downstream target, eIF2α. (c) The effect of DH20931 on the ATF4 signaling pathway and associated pro- and anti-apoptotic markers. **(C)** Densitometric analysis of the western blots from (B). Data are presented as mean ± SEM from three independent experiments. *p < 0.05, **p < 0.01, ****p < 0.0001. **(D)** The cytotoxicity of DH20931 is mediated through the ATF4 pathway. MDA-MB-468 (a), BT-474 (b), and UFH1 (c) cells expressing either a control (scrambled shRNA for a, b; pLKO1 plasmid for c) or ATF4-shRNA (shATF4) were treated with the indicated concentrations of DH20931 for 72 h. IC₅₀ values were determined by MTT cell survival assay. Data are presented as mean ± SEM from four independent estimations.

Excess lipids can alter ER membrane fluidity and composition, affecting the proper folding of newly synthesized proteins. This leads to the accumulation of misfolded or unfolded proteins in the ER lumen activating the UPR.^37^ When lipotoxic ER stress is prolonged and irreversible, the UPR shifts from a pro-survival to a pro-apoptotic program, primarily through the lipotoxic ER stress signaling axis.^21^ Upon ER stress, PERK phosphorylates eIF2α, leading to a global shutdown of protein translation but, paradoxically, the preferential translation of Activating Transcription Factor 4 (ATF4) is enhanced under these conditions. ATF4 then transcriptionally upregulates a host of pro-apoptotic genes, most notably *DDIT3*, which encodes the transcription factor CHOP (C/EBP Homologous Protein). CHOP, in turn, upregulates pro-apoptotic BH3-only proteins like PUMA, which ultimately trigger mitochondrial apoptosis.^38–40^

To investigate whether DH20931 causes lipotoxic ER stress, we analyzed the activation of the UPR pathway. Western blot analysis showed that DH20931 treatment dose-dependently increased the expression of the primary UPR stress sensors PERK, ATF6, and IRE1α (Figure 4B, panel a). Furthermore, we confirmed the specific activation of the PERK signaling cascade, observing a significant increase in phosphorylated PERK (p−PERKThr980) and its substrate, phosphorylated eIF2α (p−eIF2αSer51), in cells treated with 20 µM DH20931 for 24 h (Figure 4B, panel b). Collectively, these results demonstrate that DH20931 induces ER stress and activates the UPR.

To validate that the observed phosphorylation was a direct consequence of DH20931 activity, we employed two known PERK inhibitors, PERK inhibitor GSK2606414^41^ and the more specific inhibitor AMG PERK 44 (AMG44).^42^ As anticipated, when MDA-MB-468 cells were treated with either GSK2606414 or AMG44 alone, there was a decrease in the baseline phosphorylation levels of both PERK and eIF2α (Figure 4B, panel b, compare lane 1 with lanes 3 and 5). This confirmed the efficacy of the inhibitors in blocking this specific signaling pathway.

Co-treatment of the cells with DH20931 and PERK inhibitors revealed differential effects on the observed increase in phosphorylated PERK and eIF2α. While PERK inhibitor GSK2606414 did not affect the DH20931-induced phosphorylation, the more specific PERK inhibitor AMG44 markedly diminished it (Figure 4B, panel b, compare lane 2 with lanes 4 and 6). Total PERK and eIF2α levels remained unchanged across all treatments. These findings suggest that DH20931 activates the lipotoxic unfolded protein response pathway through PERK. The inhibition of the kinase PERK is predicted to decrease the protein abundance of its downstream target, ATF4. In agreement with this, co-treatment with the PERK inhibitors GSK2606414 or AMG44 attenuated the DH20931-induced accumulation of ATF4 protein (Figure 4B, panel b, compare lane 2 with 5 and 6). This outcome is a direct result of disrupting a critical signaling cascade that governs protein synthesis during the cellular stress response.

To further confirm that the entire PERK-eIF2α-ATF4 signaling axis was engaged by DH20931, we performed Western blot analysis on lysates from treated MDA-MB-468 cells. Treatment with DH20931 led to a significant concentration-dependent increase in the protein levels of ATF4 and its downstream targets CHOP and PUMA (Figure 4B, C). This was accompanied by a corresponding increase in the pro-apoptotic protein Bax and a decrease in the anti-apoptotic proteins Bcl-2 and Bcl-xL (Figure 4B, C). Ultimately, this signaling cascade resulted in the activation of the apoptotic machinery, evidenced by the cleavage of caspases-9 and -3, the cleavage of the caspase substrate PARP1, and the release of cytochrome c from the mitochondria into the cytosol (Figure 4B, C). Furthermore, our results demonstrated a significant, dose-dependent increase in CerS2 protein levels in response to DH20931 treatment (Figure 4B, C). These findings indicate that DH20931 likely functions through a dual mechanism. Beyond simply modulating the activity of the existing CerS2 enzyme, it also appears to augment the overall amount of enzyme available. This combined effect could yield a stronger and more lasting impact on ceramide synthesis, a crucial factor for therapeutic efficacy.

To functionally validate the essential role of the ATF4 pathway in mediating DH20931-induced cell death, we used shRNA to stably knock down ATF4 in MDA-MB-468, BT-474 and UFH1 TNBC cell lines. We then compared the sensitivity of these ATF4-knockdown cells to cells expressing a non-targeting scrambled shRNA. In all the three cell lines, the knockdown of ATF4 conferred resistance to DH20931-induced cytotoxicity (Figure 4D). This functional genetic evidence confirms that the activation of the ATF4-CHOP pro-apoptotic axis is not merely a correlative event but is a required downstream mechanism for the execution of cell death following CerS2 activation by DH20931. Interestingly, our TEM analysis also revealed a reduction in the number of mitochondria in DH20931-treated, CerS2-proficient cells (Figure 4A), suggesting that the cellular stress may also trigger mitophagy—a process of selective mitochondrial degradation—which could be a topic for future studies.

### DH20931 promotes a novel CerS2-IP3R1 interaction, enhances ER-mitochondria proximity, and triggers mitochondrial Ca²⁺ overload

The convergence of ER stress and apoptosis often occurs via the dysregulation of intracellular Ca²⁺ signaling, particularly at mitochondria-associated membranes (MAMs).^43,44^ Because CerS2 is an ER-resident enzyme and a potent inducer of ER stress, we hypothesized that it physically or functionally interacts with key Ca^2+^ regulatory proteins at the ER-mitochondria interface. We therefore focused on IP3R1, a major ER Ca^2+^ release channel highly enriched at MAMs. Using confocal immunofluorescence microscopy, we observed a strong co-localization between CerS2 and IP3R1 in 4T1-*CERS2^+/+^* cells, which increased upon treatment with DH20931 (Fig. 5A, panel a). As expected, no CerS2 signal, and thus no co-localization, was detected in the 4T1-*CERS2^−/-^* cells (Fig. 5A, panel b). Since these knockout cells were not clonally selected, a residual population of puromycin-resistant cells remained and exhibited low-level CerS2 expression (Fig. 5A, panel b, Alexa Fluor 594 channel). This is the first report of a physical interaction between a ceramide synthase and a major ER Ca²⁺ channel, revealing a previously unknown regulatory complex.

**Figure 5.**
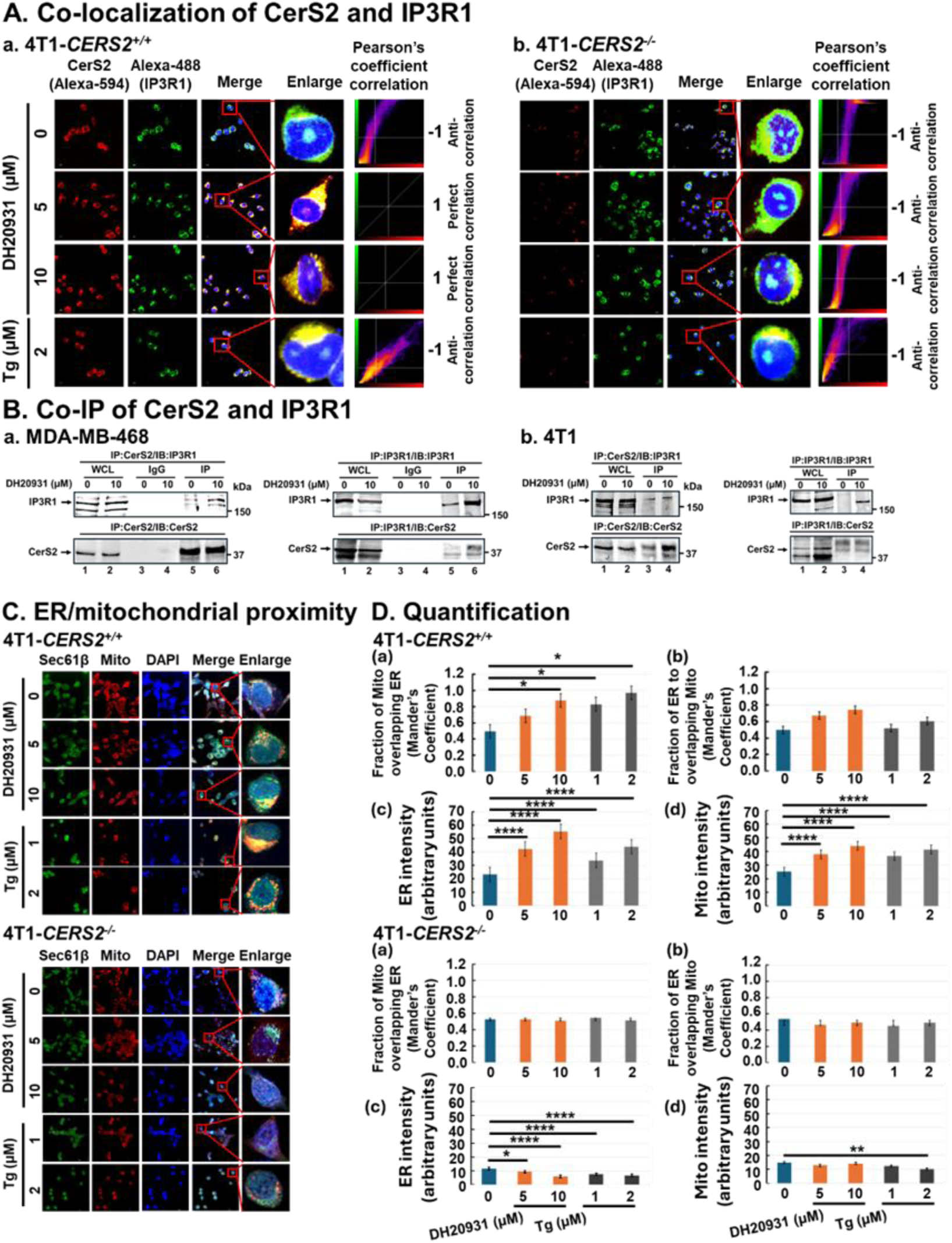
DH20931 Promotes the Interaction between CerS2 and IP3R1, Increasing ER-Mitochondria Proximity. **(A)** Immunofluorescence staining for CerS2 and IP3R1 co-localization. 4T1-*CERS2*⁺^/^⁺ (a) and 4T1-*CERS2*⁻^/^⁻ (b) cells were treated with 0, 5, or 10 µM DH20931 for 24 h. Cells were stained with anti-CerS2 (red) and anti-IP3R1 (green) antibodies. Nuclei were counterstained with DAPI (blue). Co-localization was visualized by fluorescence microscopy. **(B)** Co-immunoprecipitation (Co-IP) of CerS2 and IP3R1. Whole-cell lysates from MDA-MB468 (a) and 4T1 (b) cell lines were subjected to Co-IP using the indicated antibodies. Data are representative of three independent experiments. **(C)** Immunofluorescence staining for ER-mitochondria proximity. 4T1-*CERS2*⁺^/^⁺ and 4T1-*CERS2*⁻^/^⁻ cells were treated as in (A). Cells were stained for mitochondria (MitoTracker Red), ER (anti-Sec61β, green), and nuclei (DAPI, blue). The spatial relationship between ER and mitochondria was observed by fluorescence microscopy. **(D)** Quantification of ER-mitochondria overlap from images as in (B). Overlap was quantified using Mander’s coefficients via the JACoP plugin in ImageJ. Data are presented as mean ± SEM from 30 cells per experimental group. *p < 0.05, ****p < 0.0001.

To determine if this co-localization represented a true physical interaction, we performed co-immunoprecipitation (Co-IP) experiments. In lysates from MDA-MB468 and 4T1 cells, immunoprecipitation of endogenous IP3R1 successfully pulled down endogenous CerS2, and conversely, immunoprecipitation of CerS2 pulled down IP3R1 (Figure 5B, panels a, b). This is the first report of a physical interaction between a ceramide synthase and a major ER Ca²⁺ channel, revealing a previously unknown regulatory complex.

Since the physical tethering of the ER and mitochondria at MAMs is critical for efficient Ca²⁺ transfer^45^, we therefore investigated whether DH20931 treatment altered the proximity of these two organelles. Using confocal microscopy to visualize the ER (stained for Sec61β) and mitochondria (stained with MitoTracker Red), we observed that treatment with DH20931 significantly increased the overlap between the ER and mitochondria in 4T1-*CERS2*⁺^/^⁺ cells (Figure 5C). The quantification of this overlap using Mander’s coefficient confirmed a significant increase in ER-mitochondria proximity (Figure 5D). This effect was entirely CerS2-dependent, as no change in proximity was observed in 4T1-*CERS2*⁻^/^⁻ cells. This suggests that the CerS2-IP3R1 interaction may occur at MAMs and function to stabilize or enhance these organelles contact sites.

An enhanced flux of Ca²⁺ from the ER to the mitochondria is a functional consequence of increased ER-mitochondria proximity and potential modulation of IP3R1 channel activity.^46^ To directly measure this, we utilized Rhod-2 AM, a fluorescent indicator that specifically accumulates in mitochondria and reports on its Ca²⁺ concentration. A key finding of this study is the novel and essential role of CerS2 in mediating mitochondrial Ca²⁺ overload. Upon treatment with DH20931, 4T1-*CERS2*⁺^/^⁺ cells demonstrated a rapid increase in mitochondrial Ca²⁺ levels (Figure 6A, panels a, c). This lethal Ca²⁺ overload was completely dependent on CerS2, as it was entirely abrogated in the 4T1-*CERS2*⁻^/^⁻ cells (Figure 6A, panels b, c). A similar dose-dependent increase in mitochondrial Ca²⁺ was observed in MDA-MB-468 cells (Figure S6).

**Figure 6.**
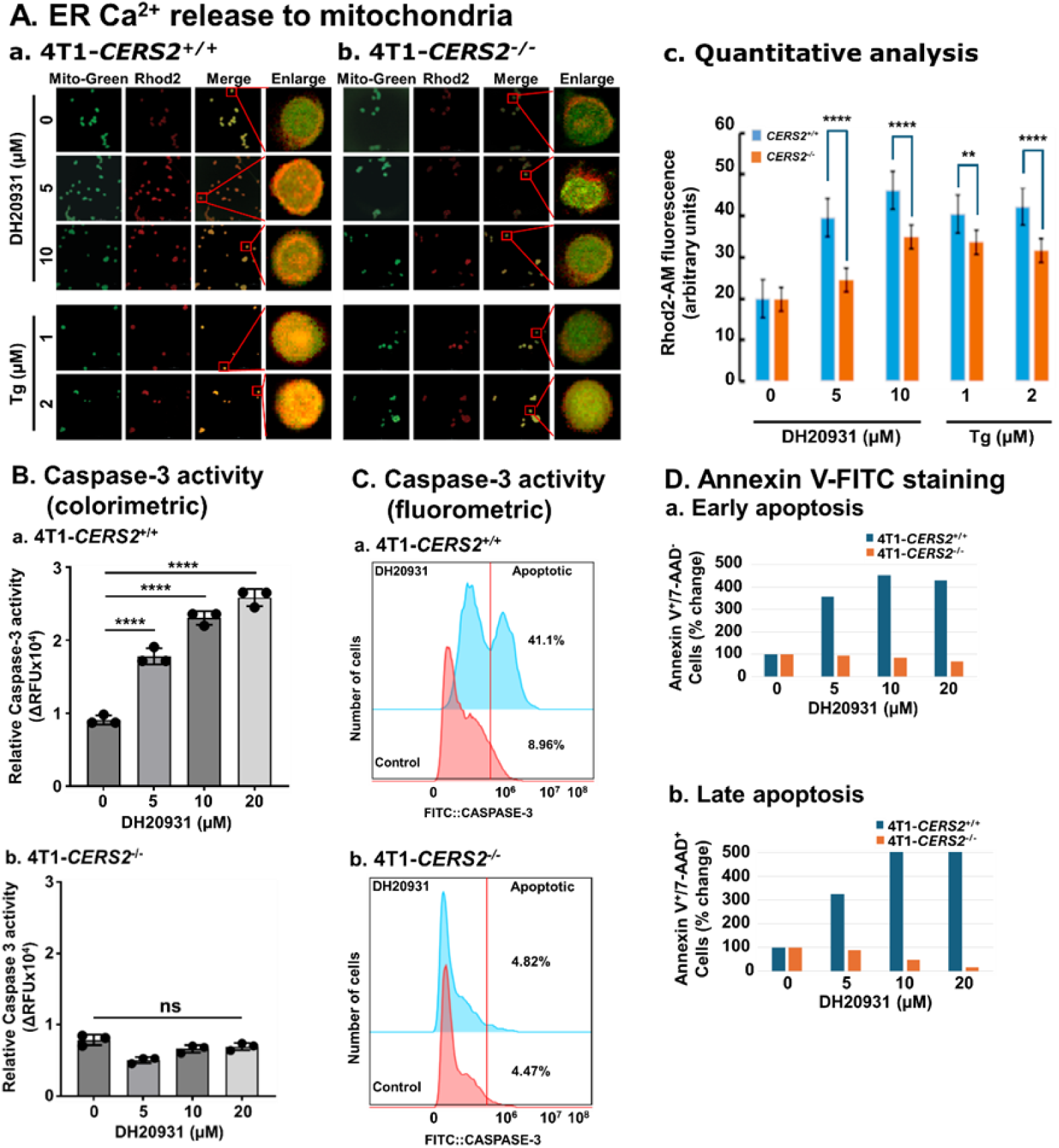
DH20931 Elevates Mitochondrial Ca²⁺ Load and Induces Apoptosis in a CerS2-Dependent Manner. **(A)** Confocal microscopy analysis of mitochondrial Ca²⁺. 4T1-*CERS2*⁺^/^⁺ (a) and 4T1-*CERS2*⁻^/^⁻ (b) cells were treated with DH20931 for 6 h and stained with Rhod-2 AM to visualize mitochondrial Ca²⁺. (c) Quantification of mitochondrial Ca²⁺ levels from fluorescence intensities. Data were measured using ImageJ and are presented as mean ± SEM from 30 cells per experimental group. ****p < 0.0001. See also Figure S6. **(B)** Caspase-3 Activity (colorimetric): 4T1-*CERS2^+/+^* (a) and 4T1-*CERS2^−/-^*(b) cells were treated with varying concentrations of DH20931 for 24 h. Caspase-3 activity was measured using the Caspase-3 activity kit. Data are presented as Mean ± SE of three independent estimations. ****p<0.0001. See also Figure S7. **(C)** Caspase-3 Activity (fluorometric): 4T1-*CERS2^+/+^* (a) and 4T1-*CERS2^−/-^*(b) cells were treated with varying concentrations of DH20931 for 24 h. Caspase-3 activity was measured using the CaspGlow^TM^ Fluorescence Active Caspase-3 Staining kit. Data are representative of two experiments. **(D)** Flow cytometric analysis of apoptosis. 4T1-*CERS2^+/+^* (a) and 4T1-*CERS2^−/-^*(b) cells were treated with varying concentrations of DH20931 for 24 h. Apoptosis was measured by APC Annexin V Staining kit. Data are normalized to 10,000 cells in each group and has been repeated twice.

Intriguingly, our findings also reveal a previously uncharacterized role for the CerS2-IP3R1 interaction in response to thapsigargin (Tg), a well-known ER stress-inducing agent. While Tg is known to inhibit SERCA pumps, leading to ER Ca²⁺ depletion and subsequent mitochondrial uptake^47,48^, our study demonstrates that this process is critically dependent on CerS2. Tg treatment resulted in an increase in mitochondrial Ca²⁺ levels, which was completely abolished in the absence of CerS2 expression (Figure 6A, panels b, c).

Previous studies have presented conflicting views on the role of IP3R1 in Tg-mediated Ca²⁺ release, with some suggesting it is independent,^49^ while others propose a modulatory role.^50^ Our research provides compelling evidence that Tg promotes the interaction between CerS2 and IP3R1, facilitating ER Ca²⁺ release and its subsequent accumulation in the mitochondria. This is substantiated by the complete blockage of Tg-induced mitochondrial Ca²⁺ accumulation in 4T1-*CERS2*^−/-^ cells (Figure 6A, panels b-c).

These results establish a clear mechanistic link, highlighting the novel role of the CerS2-IP3R1 interaction as a critical mediator of both DH20931- and Tg-induced catastrophic Ca²⁺ overload into the mitochondria—a known potent trigger of apoptosis.^51^

### DH20931 induces robust apoptosis in breast cancer cells in a CerS2-dependent manner

The culmination of severe ER stress and mitochondrial Ca²⁺ overload is the induction of programmed cell death. To confirm that DH20931-induced cytotoxicity proceeds via apoptosis, we performed a series of definitive assays. First, we measured the activity of the executioner Caspase-3, which is the final protease activated in the apoptotic cascade. Using a colorimetric substrate, we found that DH20931 induced a strong, dose-dependent increase in Caspase-3 activity in both MDA-MB-468 (Figure S6) and 4T1-*CERS2*⁺^/^⁺ cells (Figure 6B, panel a). This activation was entirely dependent on the presence of CerS2, as no increase in caspase activity was detected in the resistant 4T1-*CERS2*⁻^/^⁻ cells (Figure 6B, panel b). Similarly, using a fluorometric CaspGLOW^TM^ assay system, a CerS2-dependent Caspase-3 activation was observed in 4T1-*CERS2*⁺^/^⁺ but not in 4T1- *CERS2*⁻^/^⁻ cells-treated with 5 µM of DH20931 for 48 h (Figure 6C, panel a-b).

As a complementary approach, we employed flow cytometry to measure Annexin V staining, which identifies the externalization of phosphatidylserine—an early hallmark of apoptosis.^52^ Treatment of 4T1-*CERS2*^+/+^ cells with DH20931 for 24 h resulted in a significant and dose-dependent rise in the percentage of cells in early (Annexin V-positive) and late (Annexin V and 7-AAD-positive) apoptosis (Figure 6D, panel a). In stark contrast, DH20931 failed to induce apoptosis in 4T1-*CERS2*^−/-^ cells, as shown by the lack of increased Annexin V/7-AAD staining even at high concentrations (Figure 6D, panel b). Together, these orthogonal assays confirm that DH20931 kills breast cancer cells by triggering a robust, CerS2-dependent apoptotic pathway, thereby connecting the initial enzyme activation to the eventual demise of the cancer cell.

## DISCUSSION

In this study, we identify and characterize DH20931 as a first-in-class, small-molecule agonist of ceramide synthase 2 (CerS2), establishing it as a druggable vulnerability across diverse breast cancer subtypes. We demonstrate that pharmacological activation of CerS2 by DH20931, which shows nanomolar potency in reconstituted assays, triggers a potent, dual-pronged mechanism of apoptosis. The specificity of this action is confirmed by the resistance of *CERS2*-knockout cells to the compound, which fail to accumulate the enzyme’s very-long-chain ceramide (VLCC) products upon treatment. These findings establish that CerS2 is the critical mediator of DH20931’s anticancer activity.

By targeting a fundamental metabolic enzyme, this strategy holds promise against a wide range of tumors that exhibit altered lipid metabolism. Accordingly, DH20931 is potently cytotoxic to triple-negative (TNBC), Luminal A, Luminal B, and HER2-positive breast cancer cell lines. This broad efficacy suggests a therapeutic potential that may circumvent the resistance mechanisms that limit current hormone- and HER2-targeted therapies, establishing a new treatment paradigm for aggressive and refractory breast cancers.^53^

Our work provides a deep mechanistic understanding of how CerS2 activation drives cell death, beginning with the induction of lipotoxic endoplasmic reticulum (ER) stress. The accumulation of VLCCs, the direct products of CerS2 activity, acts as a potent insult to the ER, visualized by severe organellar dilation. This stress engages the pro-apoptotic arm of the integrated stress response (ISR), specifically activating the PERK-eIF2α-ATF4 signaling axis.^21,54^ ATF4, a master transcriptional regulator, subsequently upregulates the C/EBP Homologous Protein (CHOP), a central executioner of ISR/ER stress-induced apoptosis ^55^. The ATF4/CHOP complex then induces key death effectors, including PUMA, a potent BH3-only protein that neutralizes anti-apoptotic Bcl-2 family members at the mitochondria.^38–40^ The functional necessity of this pathway was definitively established through genetic intervention; RNAi-mediated knockdown of *ATF4* rendered breast cancer cells significantly resistant to DH20931, confirming that the PERK-ATF4-CHOP-PUMA axis of ISR is a critical mediator of the DH2091’s activity.

In parallel with ER stress, CerS2 activation engages a second, equally lethal pathway centered on the dysregulation of intracellular calcium (Ca^2+^) signaling. The ER serves as the cell’s principal Ca^2+^ reservoir, and its communication with mitochondria at specialized microdomains known as mitochondria-associated membranes (MAMs) is a critical nexus for apoptosis.^54^ While physiological Ca^2+^ transfer at MAMs is essential for bioenergetics, an uncontrolled flux of Ca^2+^ from the ER into the mitochondria is a potent and often irreversible trigger for cell death. Cancer cells frequently remodel these signaling networks to resist apoptosis, making this axis an attractive therapeutic target.^54,56^

Perhaps the most novel finding of this study is the discovery of a direct physical interaction between the metabolic enzyme CerS2 and the major ER Ca^2+^ release channel, IP3R1, which is promoted by DH20931. This discovery fundamentally reframes the biological role of CerS2, revealing it to be not merely a biosynthetic enzyme but also a regulatory or scaffolding protein that physically organizes a signaling hub at the ER-mitochondria interface. The functional consequence of this interaction is an alteration of inter-organellar communication. The DH20931-promoted CerS2-IP3R1 association correlates with a measurable decrease in the distance between the ER and mitochondria, suggesting a remodeling of MAMs into highly efficient “hotspots” for Ca^2+^ transfer.

The ultimate outcome of this reorganization is a catastrophic flux of Ca^2+^ out of the ER, through the IP3R1 channel, and directly into the mitochondrial matrix.^57^ This mitochondrial Ca^2+^ overload represents a point of no return, triggering the opening of the mitochondrial permeability transition pore (mPTP).^58^ The opening of the mPTP dissipates the mitochondrial membrane potential, halts ATP synthesis, and causes the rupture of the outer mitochondrial membrane. This leads to the release of cytochrome c, which drives the assembly of the apoptosome and the activation of the intrinsic caspase cascade, irreversibly committing the cell to apoptosis.^59^

This dual-pronged mechanism constitutes a highly effective and coordinated assault on the cancer cell. It simultaneously generates a primary “stress” signal (lipotoxic ER stress, which upregulates pro-apoptotic proteins like PUMA) while also directly “priming the executioner” (sensitizing the mitochondria for death by opening the Ca^2+^ floodgates). This synergistic “one-two punch” likely lowers the threshold for apoptosis and severely limits the cell’s opportunities for adaptation or escape, providing a powerful strategy for overcoming the inherent resistance of aggressive cancers.

To assess its clinical translatability, DH20931 was tested in a patient-derived xenograft (PDX) model of TNBC. PDX models, which retain the complex architecture and heterogeneity of the original human tumor, are considered a gold standard for preclinical evaluation. In this highly relevant and stringent model, DH20931 demonstrated remarkable efficacy, significantly inhibiting tumor growth. This success, achieved at well-tolerated doses, indicates a favorable therapeutic window and provides compelling preclinical evidence for the compound’s potential.

In conclusion, this work validates CerS2 as a bona fide druggable target in breast cancer and provides a strong rationale for advancing DH20931 toward clinical development. The robust efficacy in clinically relevant models, a promising safety profile, and novel dual mechanism of action offer a clear path forward for treating not only TNBC, for which targeted options are scarce, but also potentially for patients with Luminal or HER2-positive cancers who have developed resistance to standard-of-care therapies. The strategy of attacking the cell on two synergistic fronts may also present a higher barrier to the development of therapeutic resistance, a common downfall of therapies that target a single pathway.

### Limitations of the study

While this study establishes DH20931 as a potent CerS2 activator, several limitations warrant acknowledgment. The predicted binding interaction between DH20931 and CerS2, though supported by robust computational modeling, awaits definitive experimental validation through techniques like site-directed mutagenesis. Furthermore, while we demonstrate a novel CerS2-IP3R1 complex, the directness of this interaction and the precise mechanism by which DH20931 promotes it remain to be fully elucidated, and the compound’s selectivity against other ceramide synthase isoforms was not profiled. Our preclinical evaluation, while promising, was limited to primary tumor growth inhibition in a single PDX model and did not assess the compound’s impact on metastasis, the potential for acquired resistance, or its efficacy in combination with standard-of-care therapies. Future work should therefore focus on these areas to fully define the therapeutic potential and translational relevance of targeting CerS2 in triple-negative breast cancer.

## Acknowledgements

This work was funded by the Florida Department of Health grant 23K05-JEK to S.N. This work was funded in part by the following grants to BL: NIH/NCI R21 CA252400, NIH/NCI R21 CA277485, Florida Department of Health, James & Esther King Cancer Research Program grants 23K06 and 22K04, Florida Department of Health, Bankhead-Coley Research Pro-gram grant 23B03, and a grant from the Florida Breast Cancer Foundation. O.A.G. was supported by NIH R01 DK121831 and the Oxnard Family Foundation. B.Y. was supported by the Ocala Royal Dames for Cancer Research, Inc and the ACS-IRG to the University of Florida Health Cancer Center (UFHCC). UFHCC is an NCI-designated cancer center (P30 CA247796). The work is also partially supported by NIH grants. (CA269662, CA260239 and CA290792) and endowment funds from the Dr. and Mrs. James Robert Spenser Family to W.Z. We sincerely acknowledge the help of Mary Gragg and Karen L. Kelley at UF ICBR Electron Microscopy (RRID:SCR_019146) facility for TEM imaging. We thank Nayeong Koo for preparing and Ashwin Akki for reading the pathological slides. The authors thank the Organic Synthesis Shared Resource (OSSR) of the Penn State Cancer Institute for DH20931 synthesis and computational modeling. The graphical abstract was designed using bioRender: Scientific Image and Illustration software (biRender.com).

## Authors contributions

S.N. developed and oversaw the study, performed data analysis, acquired funding, and wrote the manuscript. H.A., H.H.N., L.A., A.S., I.M., A.G, D.B, B.Y., and C.K.M. conducted experiments and/or performed analysis. H.A. also contributed to writing the first draft of the manuscript. B.K.L., M.Z.-K., C.V., O.A.G., A.K.S., T.J.G., C.D.H., W.Z. and S.H. provided reagents, analyzed data, provided key technical and conceptual advice, and provided comments and suggestions on the final draft of the manuscript.

## Resource availability

All requests for information, data, and resources should be directed to the lead contact, Satya Narayan.

## Materials availability

All unique reagents generated in this study, including the 4T1-*CERS2*^−/-^ cell line and the DH20931 compound, are available from the Lead Contact upon completion of a Materials Transfer Agreement. Further information and requests for resources and reagents should be directed to and will be fulfilled by the Lead Contact.

## Data and code availability

The datasets generated and/or analyzed during the current study are available from the corresponding author upon reasonable request. All data supporting the findings of this study are available within the paper and its supplementary information files. The raw data for the lipidomic analysis have been deposited in the Zenodo.com under the accession code (https://zenodo.org/records/16803523).

All software used for analysis is publicly available or commercially licensed and is detailed in the Key Resources Table and STAR★Methods section. No custom code was generated for the analyses in this study. Any further information required to reanalyze the data reported in this paper is available from the Lead Contact upon request.

## STAR★METHODS

### KEY RESOURCES TABLE

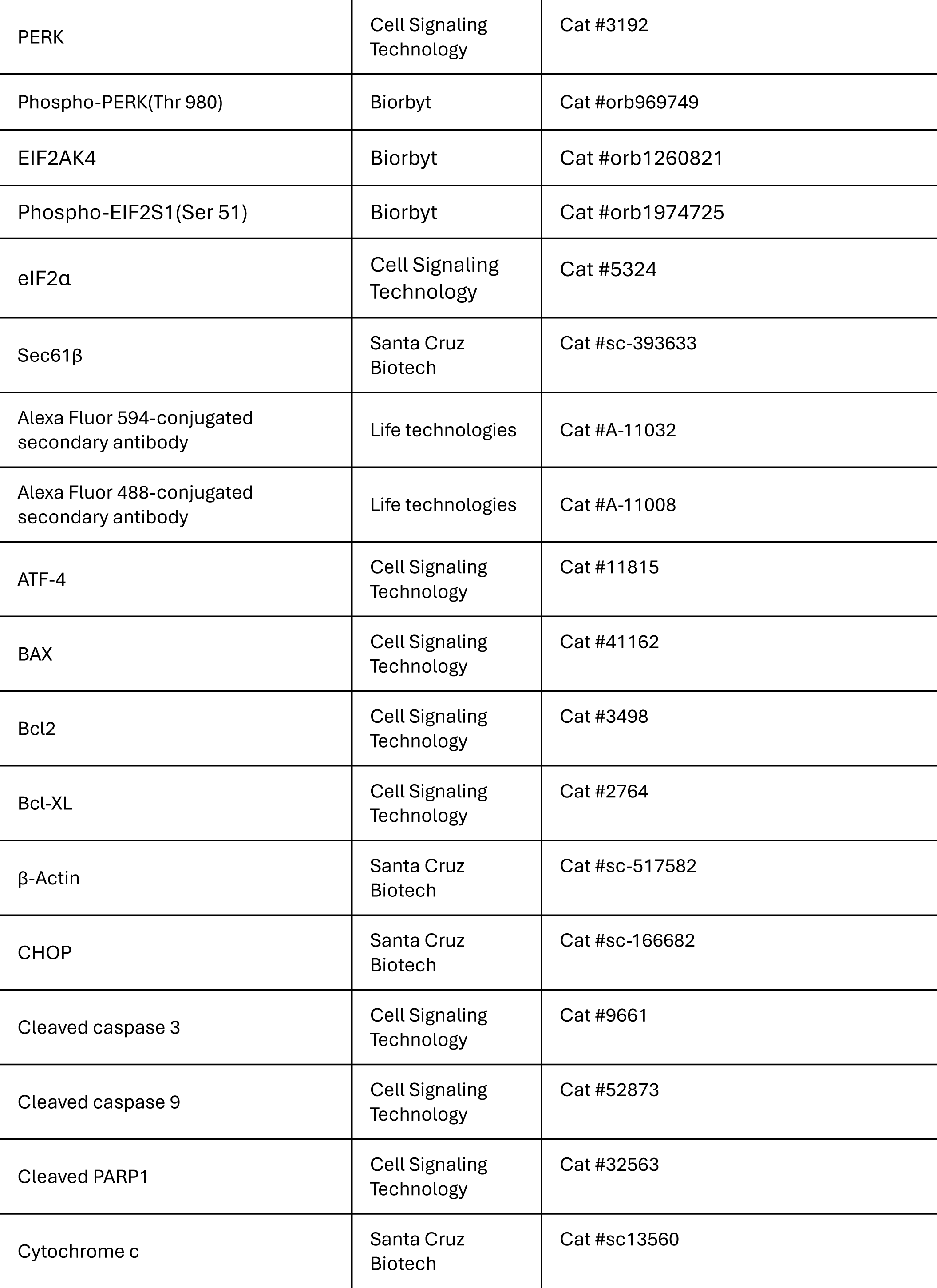
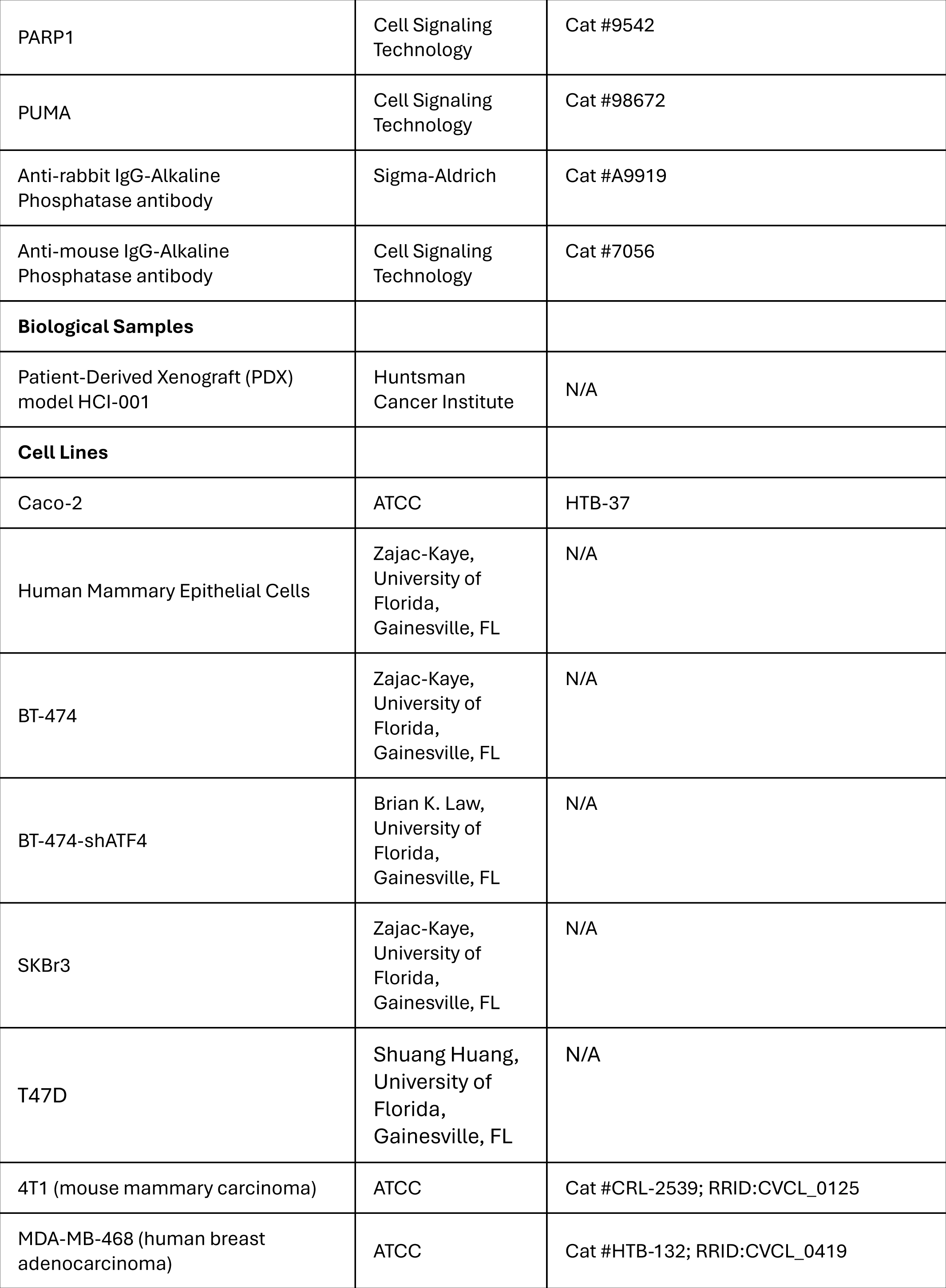
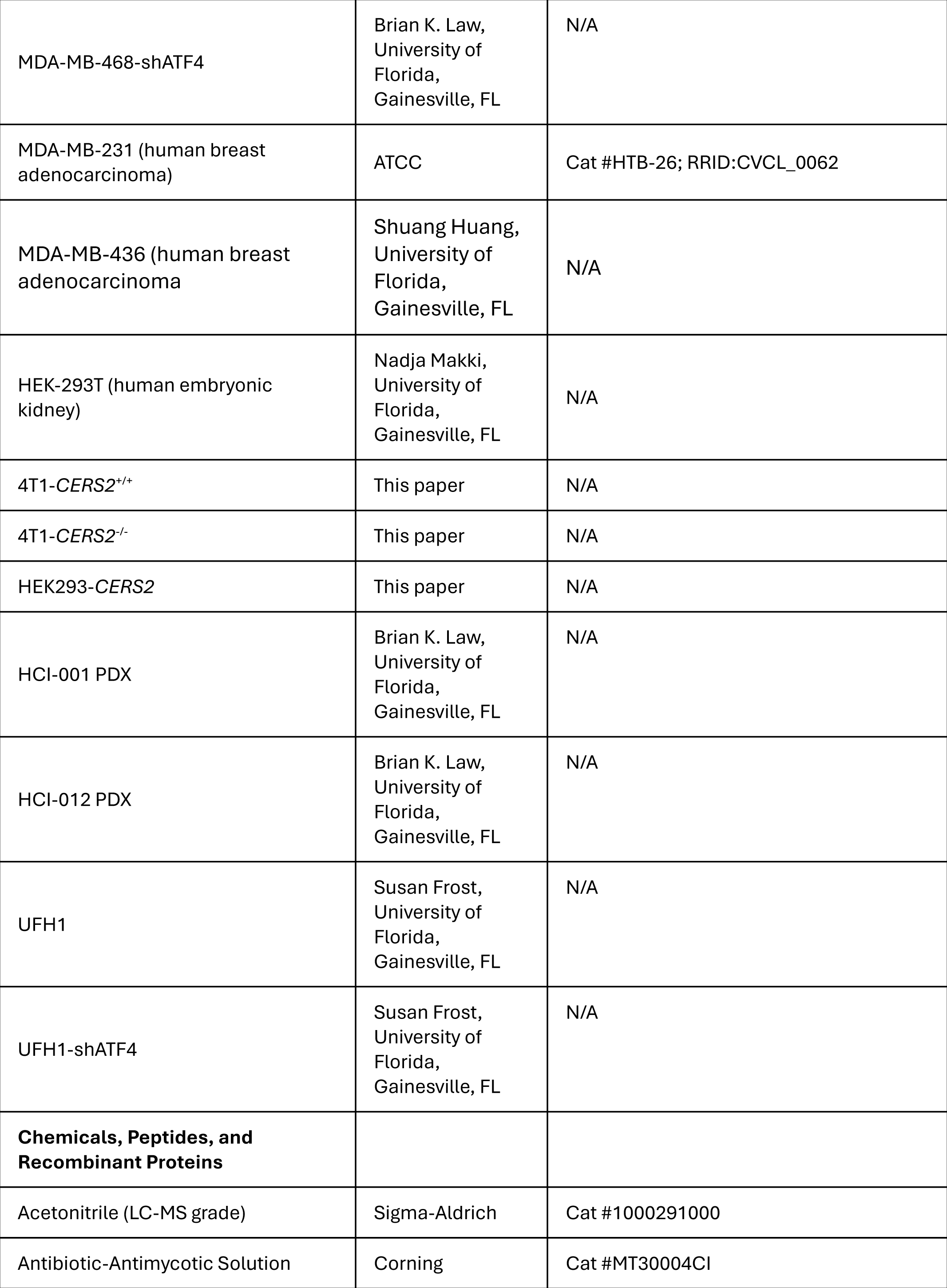
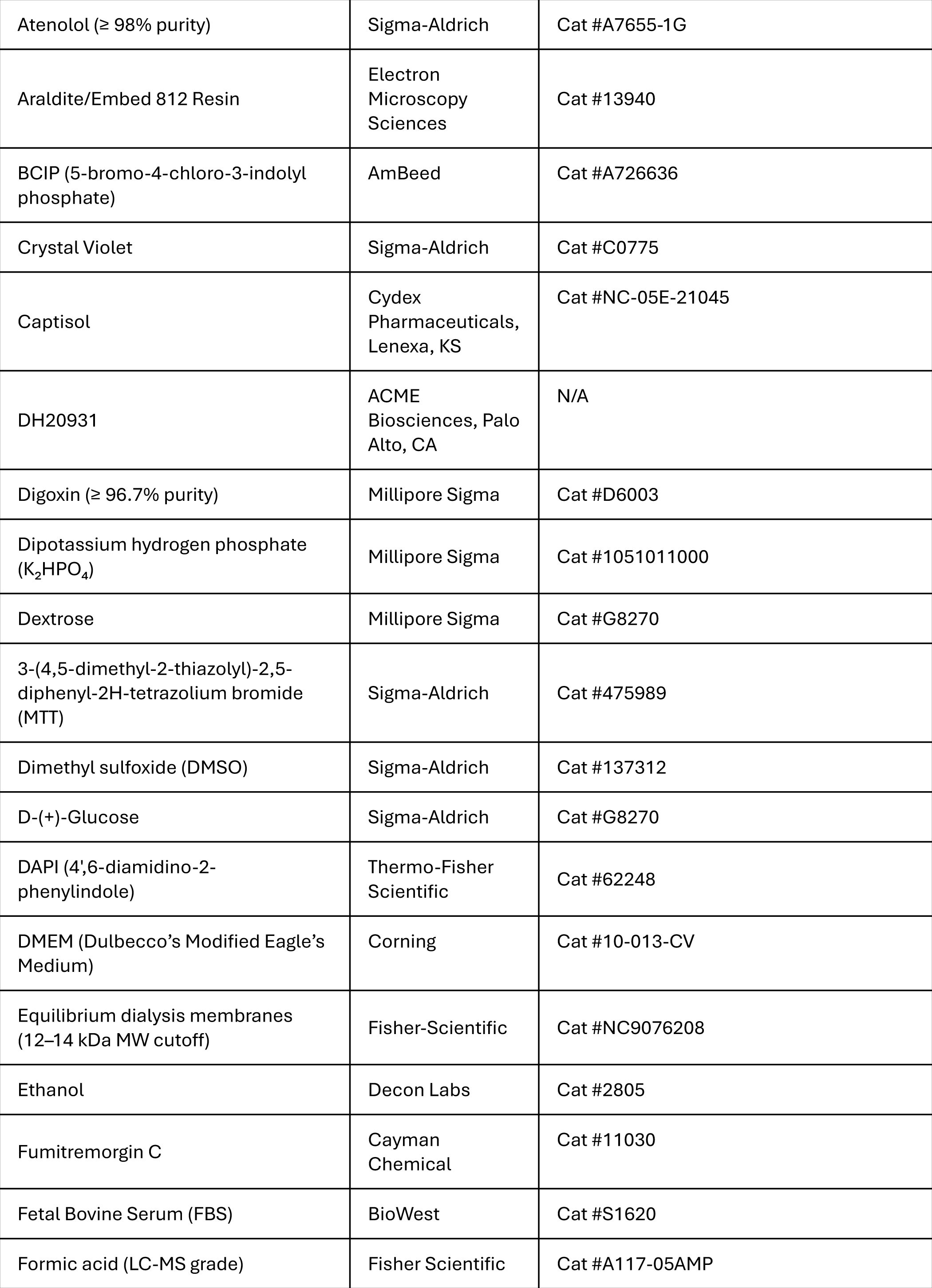
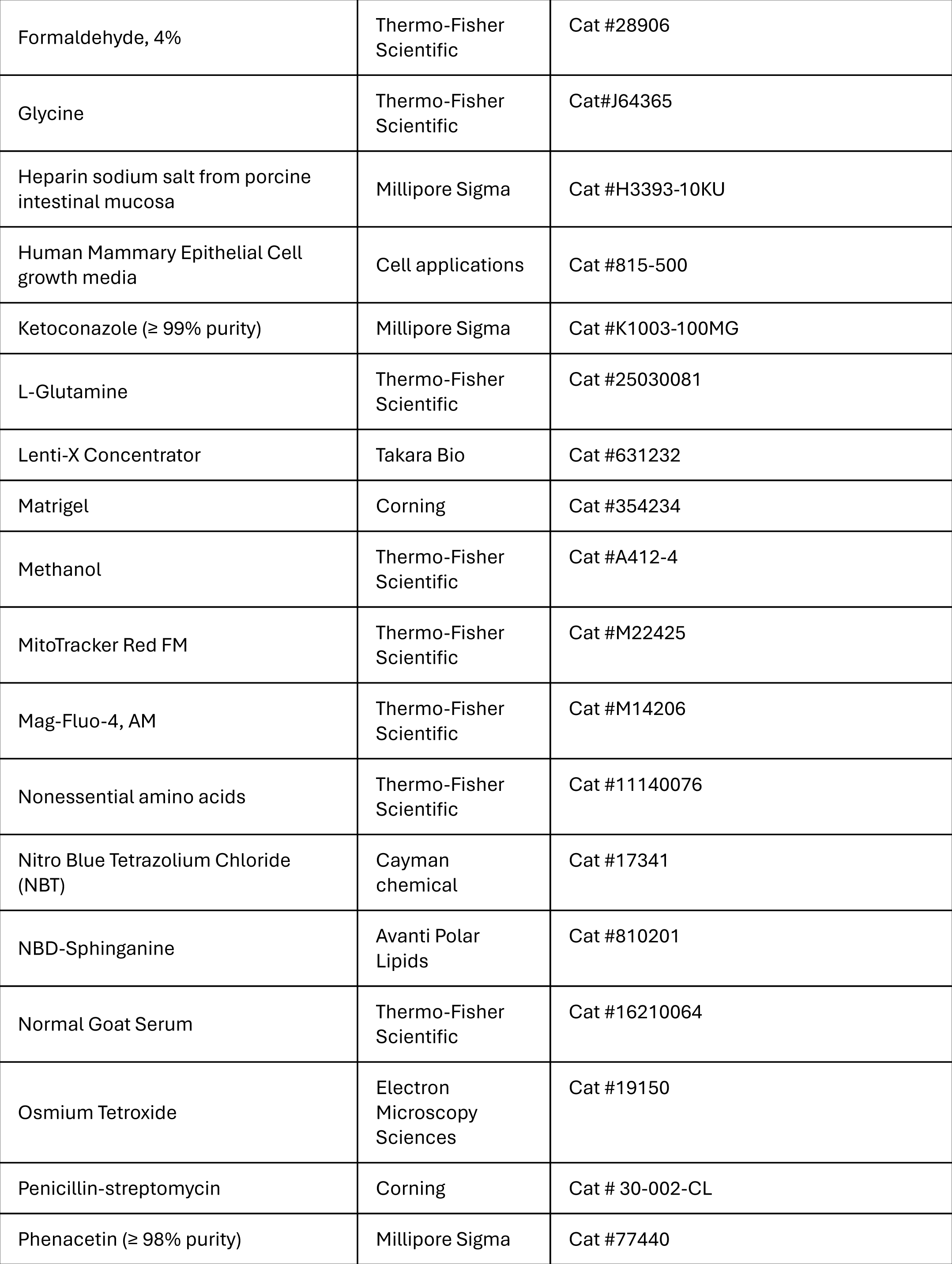
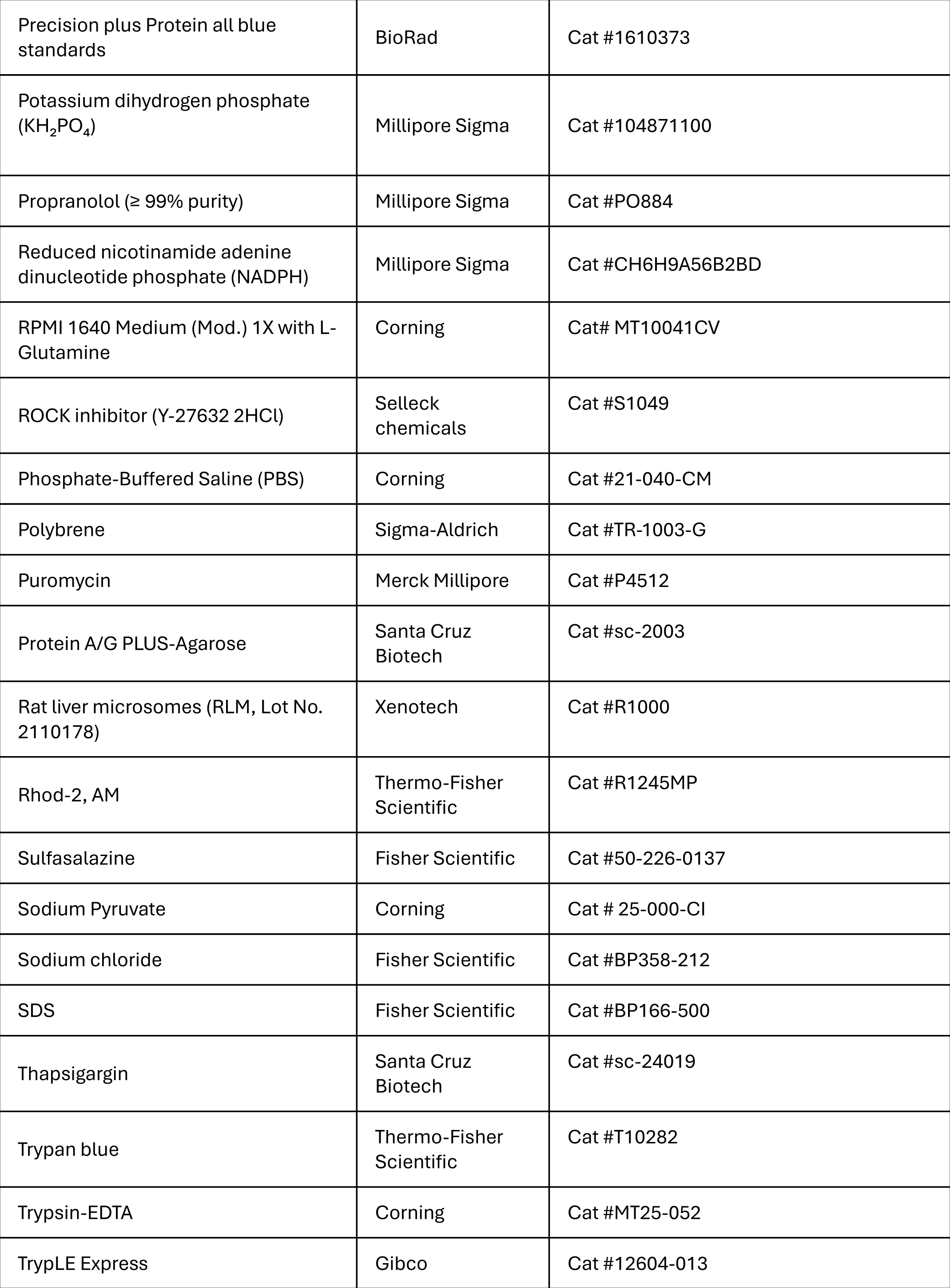
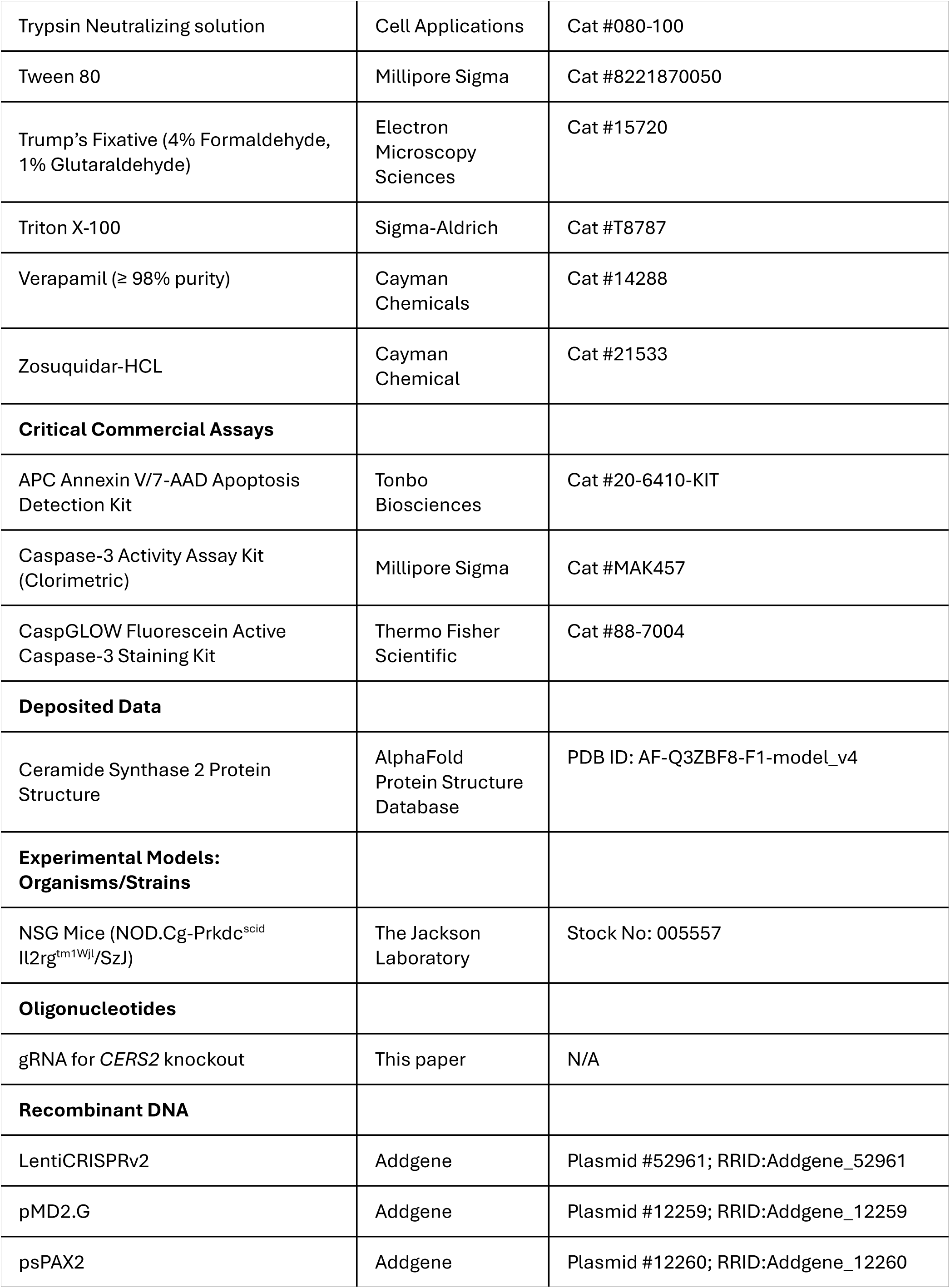
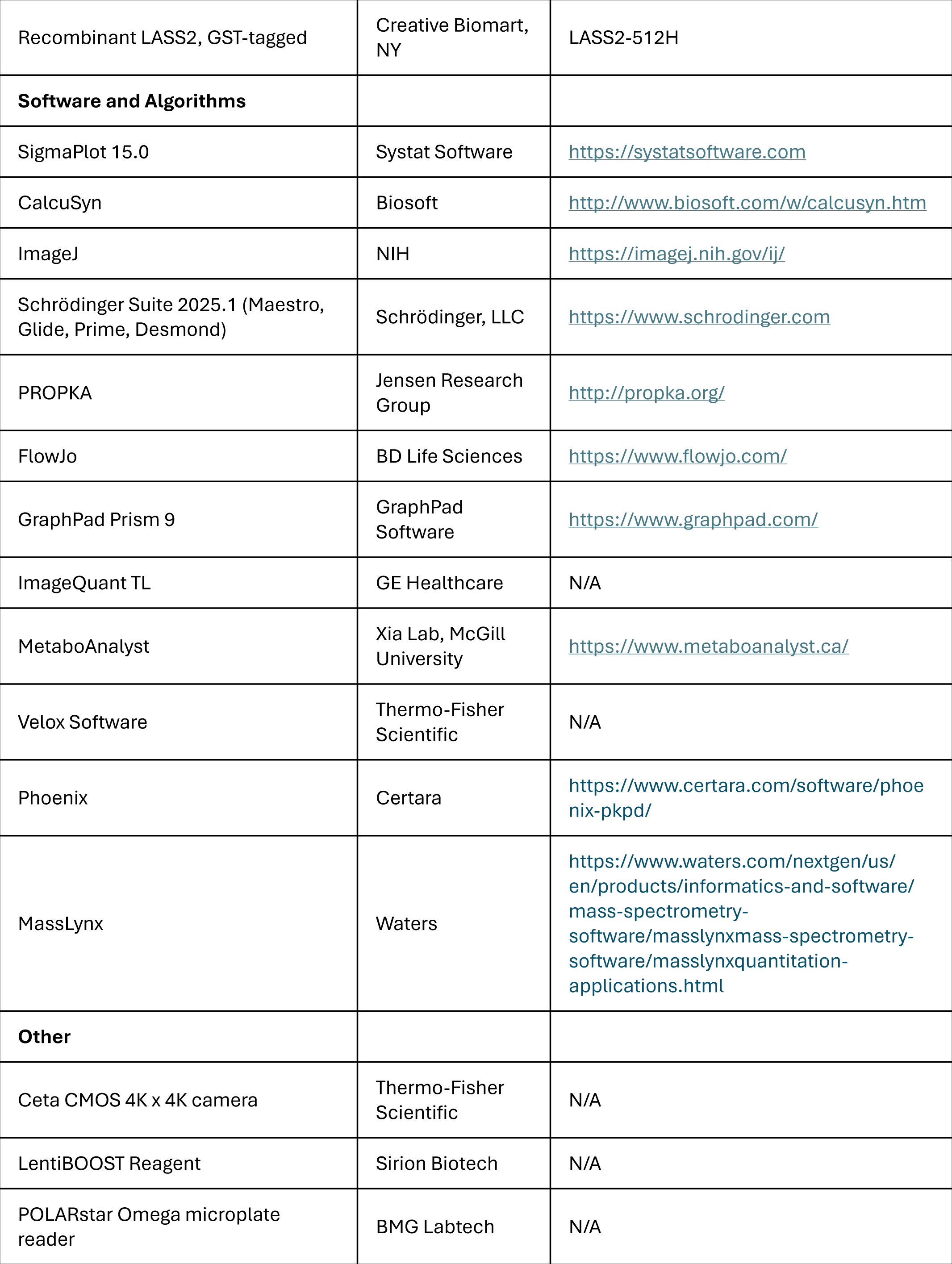
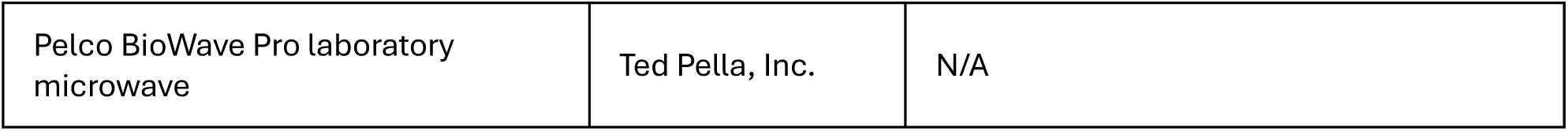

### EXPERIMENTAL MODEL AND SUBJECT DETAILS

#### Cell lines

The murine triple-negative breast cancer (TNBC) cell line 4T1 and its derivatives (4T1-*CERS2*^+/+^ and 4T1-*CERS2*^−/-^) were cultured in RPMI-1640 Medium. The human TNBC cell lines MDA-MB-468, MDA-MB-231, MDA-MB-436, T47D, BT474 and SKBr3 cell lines were cultured in Dulbecco’s Modified Eagle Medium (DMEM). All media were supplemented with 10% Fetal Bovine Serum (FBS), 2 mM L-glutamine, 100 mg/mL sodium pyruvate, and 100 U/mL penicillin-streptomycin. HEK293-*CERS2* overexpressing cells, used for activity assays, and parental HEK293 cells, used for lentivirus production, were cultured in DMEM with 10% FBS. PDX cell lines HCI-001 (TNBC) and HCI-012 (HER2+) were established in Brian Law’s lab at UF^60^ and maintained in in conditional cell reprogramming medium.^61–64^ Normal human mammary epithelial cells (HMEpC) were cultured in Human Mammary Epithelial Cell growth media containing growth factors. All cell lines were maintained at 37°C in a humidified atmosphere of 5% CO_2_. Cell lines were authenticated by STR profiling and routinely tested for mycoplasma contamination.

#### Animal models

All animal procedures were conducted in accordance with the guidelines and protocols approved by the University of Florida Institutional Animal Care and Use Committee. Female NSG (NOD.Cg-*Prkdc*^scid^ *Il2rg*^tm1Wjl^/SzJ) mice (8-10 weeks old) obtained from Jackson Laboratory (Bar Harbor, ME) were bred and housed in the specific-pathogen-free Animal Care Service (ACS) facility at the University of Florida. Mice were monitored twice weekly for tumor size and body weight. For the orthotopic xenograft model, 2 × 10⁶ MDA-MB-468 cells or 1 × 10⁶ MDA-MB-231 cells suspended in 100 µL sterile PBS (1:1, v/v with Matrigel) were injected into the left mammary fat pad. For the Patient-Derived Xenograft (PDX) model, tissue from the HCI-001 line was minced, homogenized in cold PBS, and a 100 µL suspension was injected into the left mammary fat pad ^34^.

### METHOD DETAILS

#### Cell viability assay (MTT)

The half-maximal inhibitory concentration (IC₅₀) of DH20931 was determined for human and mouse breast cancer cell lines to evaluate its effect on cell proliferation. The murine 4T1-*CERS2*^+/+^ and 4T1-*CERS2*^−/-^ cell lines were cultured in RPMI-1640 medium, while the human MDA-MB-468 cell line was grown in DMEM. All culture media were supplemented with 2 mM glutamine, 100 mg/ml sodium pyruvate, 4.5 g/L glucose, and 10% Fetal Bovine Serum (FBS). The cells were maintained in a humidified incubator at 37°C with a 5% CO₂ atmosphere.

Cells were seeded into 96-well plates at a density of 2 × 10^3^ cells per well and allowed to adhere overnight. Following attachment, the cells were treated with DH20931 at concentrations (0, 0.5, 1, 2.5, 5, 7.5, 10, 25, and 50 µM). After a 72-hour incubation period, cell viability was assessed using the MTT [3-(4,5-Dimethyl-2-thiazolyl)-2,5-diphenyl-2H-tetrazolium Bromide] assay, as described earlier.^65^ A 10 µL volume of MTT solution (5 mg/mL in PBS) was added to each well, and the plates were incubated for an additional 4 h. During this time, viable cells metabolize the yellow MTT tetrazolium salt into purple formazan crystals. To solubilize the formazan crystals, 100 μL of dimethyl sulfoxide (DMSO) was added to each well. The plates were then left overnight at room temperature to ensure complete dissolution.

The absorbance of each well was measured spectrophotometrically at a wavelength of 570 nm using a POLARstar Omega microplate reader. The resulting data were used to generate dose-response curves by plotting cell viability against the logarithm of the drug concentration. IC₅₀ values, representing the concentration of DH20931 required to inhibit cell proliferation by 50%, were calculated from these curves. The results are presented as the mean ± standard error from three independent experiments and were analyzed using SigmaPlot15 software.

#### 3D on-top spheroid formation and drug treatment assay

To prepare for the three-dimensional (3D) cultures, the bottoms of 48-well clear-bottom plates were coated with a base layer of Matrigel. A stock solution of Matrigel was diluted 1:1 (v/v) in phenol red-free medium. Sixty microliters (60 µL) of this diluted Matrigel were added to each well. The plates were then incubated at room temperature for 10 min, followed by 30 min in a humidified incubator at 37°C and 5% CO₂ to allow the Matrigel to solidify.

Cell suspensions of MDA-MB-468 and BT-474 were prepared by mixing the cells in culture medium with an equal volume of the diluted Matrigel solution (1:1 v/v). Specifically, 500 MDA-MB-468 cells or 1,000 BT-474 cells were suspended in a final volume of 120 µL. This cell-Matrigel suspension was then carefully pipetted on top of the pre-solidified Matrigel base layer in each well. The plate was returned to the incubator for 1 h at 37°C to allow the top layer to gel and to encourage the initiation of spheroid formation. Following this, 250 µL of complete culture medium was gently added to each well. The cells were then incubated for 3 days.

On day 3, the existing medium was carefully replaced with fresh medium containing DH20931 at final concentrations of 0, 5, or 10 µM. The spheroids were then cultured for an additional 7 days. On day 10, images of spheroids were captured using Leica DM4000B microscope at 5X magnification to assess their growth and morphology. The experiment was performed in duplicate, and four distinct images were taken from each well for subsequent analysis and spheroid counting.

#### Generation of 4T1 knockout cell lines via CRISPR-Cas9

To generate gene-specific knockouts in 4T1 cells, guide RNAs (gRNAs) were cloned into the LentiCRISPRv2 lentiviral vector. The cloning was performed using the Golden Gate assembly strategy as previously described.^66,67^ Lentiviral particles were produced by co-transfecting the LentiCRISPRv2 constructs, each harboring a specific gRNA, into 293T cells along with the envelope plasmid pMD2.G and the packaging plasmid psPAX2.

The day before transfection, HEK-293T cells were seeded in T25 cm² flasks. One hour prior to the transfection, the culture medium was replaced with 2.25 mL of Opti-MEM medium. A DNA mixture was prepared containing 3.4 µg of the lentiCRISPRv2-gRNA plasmid, 1.7 µg of pMD2.G, and 2.6 µg of psPAX2 diluted in 700 µL of Opti-MEM with 35 µL of PLUS™ Reagent. Concurrently, a lipid mixture was prepared by diluting 17.25 µL of Lipofectamine 2000 in 700 µL of Opti-MEM. Both mixtures were incubated for 5 minutes at room temperature before being combined and incubated for an additional 20 min to allow for the formation of lipofectamine-DNA complexes. This complex was then added to the 293T cells and were incubated for 6 h at 37°C and 5% CO₂.

After the 6-hour transfection, the medium was replaced with 6 mL of a specialized lentivirus harvest medium (DMEM high glucose, 10% FBS, 1X Penicillin/Streptomycin, and 1% BSA). At 60 h post-transfection, the virus-containing supernatant was collected and centrifuged at 3000 rpm for 10 min at 4°C to pellet cell debris. The clarified supernatant was then passed through a sterile 0.45 µm PES syringe filter.

To concentrate the viral particles, one-third of the supernatant volume of Lenti-X™ Concentrator was added. The mixture was gently inverted, then incubated on ice or at 4°C for 2.5 h to overnight. The lentivirus was pelleted by centrifugation at 1500 × g for 45 min at 4°C. The supernatant was discarded, and the viral pellet was resuspended in 300 µL of harvest medium, aliquoted into sterile cryovials, and stored at -80°C.

The lentiviral preparations generated were used to transduce fresh 4T1 cells via the spinoculation method. On Day 0, 0.5 × 10⁶ 4T1 cells were seeded per well in a 12-well plate. On Day 1, the centrifuge was pre-warmed to 33°C. The culture medium was replaced with 0.5 mL of fresh medium supplemented with Polybrene (8 µg/mL) to enhance transduction efficiency. For cells known to be difficult to transduce, 5 µL of LentiBOOST™ reagent per 500 µL of medium was also included. Ten microliters (10 µL) of concentrated lentivirus were added to each well, leaving some wells un-transduced to serve as controls for selection. The plate was sealed with sterile adhesive film and centrifuged at 1000 × g for 2 h at 33°C. Following centrifugation, the film was removed inside a biosafety cabinet, and the cells were incubated overnight.

Post incubation, the cells were expanded, and the clonal selection was initiated by replacing the medium with complete medium supplemented with puromycin. The specific concentration of puromycin was predetermined using a puromycin kill curve on parental 4T1 cells. The selection medium was refreshed as needed until all cells in the un-transduced control flask were killed. Once selection was complete, the medium was replaced with fresh, antibiotic-free complete medium to allow the surviving population of knockout cells to recover and expand.

#### Clonogenic survival assay

To evaluate the long-term effects of DH20931 on cell survival and proliferation, a clonogenic assay was performed using 4T1-*CERS2*^+/+^ and 4T1-*CERS2*^−/-^ cells. Cells were seeded in triplicate into 6-well plates at a low density of 500 cells per well, each containing 2.5 mL of complete RPMI growth medium. The plates were incubated for 24 h to allow the cells to attach. Following attachment, the medium was replaced with fresh medium containing DH20931 at final concentrations of 2.5, 5, or 10 µM.

The cells were incubated undisturbed for seven days to allow for the formation of distinct colonies. After the incubation period, the medium was removed, and the colonies were fixed and stained in a single step using a 0.5% crystal violet solution in 25% methanol for one hour. Following staining, the plates were washed twice with water to remove excess stain and left to air dry completely.

The dried plates were scanned to create digital images for analysis. The number of colonies, defined as clusters containing at least 50 cells, was quantified using ImageJ software. The surviving fraction for each treatment was calculated by dividing the number of colonies in the treated wells by the number of colonies in the control (untreated) wells. The data were then plotted using SigmaPlot 15 for graphical representation.

#### Western blot analysis

To examine the effect of DH20931 treatment on the expression of various proteins, cells were treated with DH20931 at concentrations of 0, 2.5, 5, 10, and 20 µM for 24 h. Western blot analysis was carried out using 5-bromo-4-chloro-3-indolyl phosphate (BCIP)/nitro blue tetrazolium (NBT) substrate as described earlier.^65^ The resulting protein bands were imaged, and their intensities were quantified using ImageJ software. The expression levels of target proteins were normalized to a loading control (e.g., β-actin) to ensure accurate comparison across samples.

#### CerS2 activity assay

HEK-293T cells were engineered to overexpress ceramide synthase 2 (*CERS2*) as described previously.^68^ The cells were transfected with a LASS2-512H plasmid, which encodes recombinant human *LASS2* (*CERS2*) with a Glutathione S-transferase (GST) tag. The transfection was performed using Lipofectamine 2000 according to the manufacturer’s protocol.

For lysate preparation, transfected HEK-293T cells from two confluent 100 mm dishes were collected. The cells were lysed in a buffer containing 20 mM HEPES (pH 7.4), 25 mM KCl, 2 mM MgCl₂, and 250 mM sucrose. To prevent protein degradation, the lysis buffer was freshly supplemented with a protease inhibitor cocktail consisting of 1 mM PMSF, 80 µg/mL leupeptin, 80 µg/mL pepstatin A, and 80 µg/mL aprotinin.

CerS2 activity in the cell lysates was quantified using a commercially available kit (Avanti Polar Lipids, Inc.) following the provided protocol.^31^ In brief, 25 µg of total protein from the cell lysate was pre-incubated with varying concentrations of the activator DH20931 (0, 0.5, 1, 2.5, and 5 µM) for 5 min. The enzymatic reaction was initiated by adding a mixture containing HEPES buffer, KCl, MgCl₂, dithiothreitol (DTT), bovine serum albumin (BSA), the fluorescent substrate NBD-sphinganine, and a CerS2-specific fatty acid-CoA (C24:0-CoA). The reaction mixtures were incubated for 1 h at 37°C.

Following the incubation, total lipids were extracted and separated by thin-layer chromatography (TLC) using a chloroform:methanol:water (8:1:0.1, v/v/v) solvent system. The fluorescently labeled product (NBD-C24:0-ceramide) was visualized, and the fluorescence intensity was quantified using an ImageQuant LAS 4000 imager. CerS2 enzymatic activity was assessed as directly proportional to the signal intensity.

#### Lipidomic analysis

Triplicate cultures of 4T1-*CERS2*^+/+^ and 4T1-*CERS2*^−/-^ cells were seeded at a density of 10 × 10⁶ cells per plate. After allowing the cells to adhere, they were treated with 5 µM DH29931 for 24 h. Following treatment, the cells were scraped from the plates and washed with ice-cold phosphate-buffered saline (PBS) to remove any residual media. The cell pellets were collected by centrifugation. To account for any variations in cell number between samples, the total protein content of a small aliquot from each pellet was determined using a bicinchoninic acid (BCA) assay. Lipid concentrations were subsequently normalized to the total protein measurement.

Lipids were extracted from the harvested cell pellets using a modified Bligh and Dyer method. This procedure involves the addition of a chloroform:methanol mixture to the cell pellet, followed by vortexing to disrupt the cells and solubilize the lipids. The addition of water induces a phase separation, resulting in an upper aqueous phase and a lower organic phase containing the lipids. The lower organic phase was carefully collected for subsequent analysis. To ensure accurate quantification and account for extraction efficiency, a suite of deuterated internal standards, representing various lipid classes, was included in the extraction solvent.

The dried lipid extracts were reconstituted and subjected to high-performance liquid chromatography (HPLC) for separation. A reverse-phase column was employed to separate the lipid species based on their hydrophobicity. The eluent from the HPLC was directly introduced into a mass spectrometer equipped with an electrospray ionization (ESI) source for sensitive detection and identification of the lipid species.

The raw mass spectrometry data files were processed using MetaboAnalyst software. This involved several key steps: peak detection, alignment of retention times across all samples, and integration of the peak areas. The abundance of each identified lipid was normalized to the corresponding internal standard for its lipid class and expressed as a relative concentration. To visualize the overall differences in the lipid profiles between the experimental groups, principal component analysis (PCA) was performed. To identify statistically significant changes in the abundance of individual lipid species between the control and DH29931-treated groups for both the 4T1-*CERS2*^+/+^ and 4T1-*CERS2*^−/-^ cell lines, unpaired t-tests were conducted.

#### Transmission Electron Microscopy (TEM)

Following experimental treatments, 4T1-*CERS2*^+/+^, 4T1-*CERS2*^−/-^, and MDA-MB-468 cell suspensions were fixed in Trump’s Fixative (Electron Microscopy Sciences) to preserve cellular structures for electron microscopy. All subsequent processing steps were performed using a Pelco BioWave Pro laboratory microwave system (Ted Pella, Redding, CA) to enhance reagent infiltration and reduce processing times. Following initial fixation, the cells were washed in buffer, pelleted by centrifugation, and encapsulated in buffered 3% low-temperature gelling agarose to form solid pellets for easier handling.

The agarose-encapsulated cell pellets were post-fixed in a buffered 2% osmium tetroxide solution to enhance lipid and membrane contrast. After a water wash, the samples were subjected to a graded ethanol dehydration series, starting from 25% and increasing in 5-10% increments to 100% ethanol, followed by a final dehydration step in 100% anhydrous acetone. The dehydrated pellets were then infiltrated with Araldite/Embed epoxy resin (Electron Microscopy Sciences) mixed with Z6040 embedding primer (Electron Microscopy Sciences). This infiltration was performed in a stepwise manner, beginning with a 3:1 ratio of anhydrous acetone to resin, followed by 1:1 and 1:3 ratios, and concluding with three changes of 100% Araldite/Embed resin to ensure complete substitution of acetone with the embedding medium.

Once the resin was polymerized, ultrathin sections of approximately 100 nm were cut from the sample blocks. These sections were collected on carbon-coated Formvar 2 x 1 mm slot copper grids (Electron Microscopy Sciences). For enhanced contrast, the sections were post-stained with 2% aqueous uranyl acetate followed by lead citrate. The prepared grids were examined using a G2 Talos L120C Transmission Electron Microscope operated at an accelerating voltage of 120 kV. Digital images of the cellular ultrastructure were acquired using a Ceta CMOS 4K × 4K camera controlled by Velox Software.

#### Immunofluorescence for CerS2 and IP3R1 co-localization

4T1-*CERS2*^+/+^ and 4T1-*CERS2*^−/-^ cells were seeded at a density of 4 × 10⁴ cells per well into 6-well plates containing 1.5 mm glass coverslips. The cells were incubated overnight to allow for attachment. Subsequently, they were treated for 24 h with either DH20931 (at 5 and 10 µM) or thapsigargin (at 1 and 2 µM) as a positive control.

Following treatment, the cells were processed for dual-label immunofluorescence as follows: Cells were gently washed with phosphate-buffered saline (PBS), then fixed with 4% formaldehyde in PBS for 15 min at room temperature. After two washes with PBS, cells were permeabilized with 0.2% Triton™ X-100 in PBS for 10 min. To prevent non-specific antibody binding, cells were incubated in a blocking buffer (10% normal goat serum in PBS) for 1 h at room temperature.

The cells were then co-incubated with a cocktail of two primary antibodies diluted in blocking buffer for 1 h at room temperature. The cocktail contained: mouse anti-CerS2/LASS2 at a 1:300 (v/v) dilution, and rabbit anti-IP3R1 at a 1:300 (v/v) dilution. After washing the cells three times with PBS, they were co-incubated for 1 h at room temperature with a cocktail of fluorophore-conjugated secondary antibodies diluted 1:500 (v/v) in blocking buffer. The plate was covered with foil to protect the samples from light. The cocktail contained: goat anti-Mouse IgG (H+L), Alexa Fluor™ 594, and goat anti-Rabbit IgG (H+L), Alexa Fluor™ 488.

Following two final washes with PBS, the coverslips were carefully mounted onto glass microscope slides using ProLong™ Diamond Antifade Mountant with DAPI for nuclear counterstaining. The coverslips were visualized using a fluorescence microscope (e.g., Nikon A1 or similar) equipped for multi-channel imaging. Images were captured using appropriate filter sets for each fluorophore: DAPI (Nuclei) (excitation ∼360 nm / Emission ∼460 nm), Alexa Fluor 488 (IP3R1) (excitation ∼495 nm / Emission ∼519 nm), and Alexa Fluor 594 (CerS2) (excitation ∼590 nm / Emission ∼618 nm).

The captured images were analyzed to assess the degree of co-localization between the CerS2 (red channel) and IP3R1 (green channel) signals. Appropriate controls, including secondary antibody-only samples, were used to confirm signal specificity.

#### ER-Mitochondria Proximity Assay

4T1-*CERS2*^+/+^ and 4T1-*CERS2*^−/-^ cells were seeded into Nunc™ Glass Bottom Dishes and cultured overnight to reach approximately 70% confluence. The experiment was performed in triplicate. Cells were then treated for 24 h with either DH20931 (at 5 and 10 µM) or thapsigargin at concentrations of 1 and 2 µM. A sequential staining protocol was used to label mitochondria in live cells followed by immunostaining for the endoplasmic reticulum (ER) in fixed cells.

Cells were first washed with Hank’s Balanced Salt Solution (HBSS). Mitochondria were then stained by incubating the cells with 100 nM MitoTracker™ Red FM in pre-warmed HBSS for 25 min at 37°C. After staining, cells were washed three times with HBSS and left to recover in fresh medium for 30 min.

Cells were fixed with 4% formaldehyde in PBS for 30 min at room temperature, followed by permeabilization with 0.2% Triton™ X-100 in PBS for 10 min. Non-specific antibody binding sites were blocked by incubating the cells with 10% normal goat serum in PBS for 1 h at room temperature.

For ER staining, cells were incubated with a primary antibody against the ER marker Sec61β (Rabbit anti-Sec61β, Abcam, Cat. No. ab78276) at a 1:300 (v/v) dilution for 1.5 h. After washing three times with PBS, cells were incubated for 1 h with a fluorophore-conjugated secondary antibody, Alexa Fluor™ 488 Goat anti-Rabbit IgG (H+L) at a 1:500 dilution. After a final wash, the coverslips were mounted onto glass slides using ProLong™ Diamond Antifade Mountant with DAPI for nuclear counterstaining.

Images were acquired using a Nikon A1R confocal microscope with a Plan APO VC 40x oil immersion objective, controlled by NIS-Elements software (Nikon Instruments Inc.). Fluorescence was detected using the following settings: Alexa Fluor 488 (ER) (excitation at 490 nm / Emission collected at 517 nm) and MitoTracker Red FM (Mitochondria) (excitation at 581 nm / Emission collected at 644 nm).

The degree of ER-Mitochondria proximity was quantified from the captured images. The extent of overlap between the green (ER) and red (mitochondria) channels was calculated by determining Mander’s overlap coefficients using the JACoP (Just Another Co-localization Plugin) for ImageJ software.

#### Measurement of ER and Mitochondrial Ca²⁺ by Confocal Microscopy

MDA-MB-468, 4T1-*CERS2*^+/+^, and 4T1-*CERS2*^−/-^ cells were seeded into Nunc™ Glass Bottom Dishes and cultured overnight to reach approximately 70% confluence. The experiment was performed in triplicate. Cells were treated for 6 h with either DH20931 (at 5 and 10 µM) or thapsigargin (at 1 and 2 µM) as a positive control for ER calcium depletion. Following treatment, cells were rinsed with Hank’s Balanced Salt Solution (HBSS). They were then co-incubated for 20 min at 37°C in a pre-warmed staining solution containing two calcium indicators: 1 µM Mag-Fluo-4, AM to report endoplasmic reticulum (ER) Ca²⁺ levels, and 100 nM Rhod-2, AM to report mitochondrial Ca²⁺ levels. After staining, the cells were rinsed three times with probe-free HBSS and then fixed with 4% formaldehyde for 15 min at 37°C. Finally, the cells were washed twice with PBS to prepare for imaging.

Fixed cells were imaged on a Nikon A1R-MPsi-STORM 4.0 confocal microscope using a 40x Plan Apochromat oil-immersion objective (NA 1.40). Fluorescence intensity was measured using the following settings: Mag-Fluo-4 (ER Ca²⁺) (excitation at 490 nm, emission collected at 517 nm) and Rhod-2 (mitochondrial Ca²⁺) (excitation at 552 nm, emission collected at 575 nm). The fluorescence intensities corresponding to ER and mitochondrial calcium levels were quantified from the captured images using ImageJ software.

#### Caspase-3 activity assay

##### Colorimetric assay

To quantify apoptosis, the enzymatic activity of Caspase-3 was measured in MDA-MB-468, 4T1-*CERS2*^+/+^ and 4T1-*CERS2*^−/−^ cell lines. Cells were treated with 10 µM DH20931 for 24 h prior to the assay. The Caspase-3 activity was assessed using a commercial fluorometric kit according to the manufacturer’s instructions. Briefly, after treatment, cells were harvested and lysed using the buffer provided in the kit. The lysates were agitated for 30 min at 4°C and then clarified by centrifugation at 2,500 × g for 10 min at 4°C to pellet cellular debris.

For the assay, 50 µL of the resulting supernatant (cell lysate) from each sample was transferred to a well on a black, flat-bottom 96-well plate. The reaction was initiated by adding 100 µL of the Caspase-3 fluorogenic working reagent to each well. The plate was then incubated for 60 min at 37°C, protected from light. The fluorescence intensity, which is directly proportional to Caspase-3 activity, was measured using a microplate reader set to an excitation wavelength of 400 nm and an emission wavelength of 490 nm. All data were subsequently analyzed using GraphPad Prism.

##### Fluorometric assay

We also determined Caspase-3 activity using the CaspGLOW™ Fluorescein Active Caspase-3 Staining Kit. 4T1-*CERS2^+/+^* and 4T1-*CERS2^−/-^* cells were seeded and treated with 5 μM DH20931 for 48 h, alongside untreated controls. Following treatment, 1 × 10^6^ cells/mL were incubated with 1 μL FITC-DEVD-FMK at 37°C for 60 min in a humidified 5% CO_2_ incubator. Where indicated, Z-VAD-FMK (1 μL/mL) was added as a caspase inhibitor control to confirm staining specificity. After incubation, cells were washed twice with the provided wash buffer, centrifuging at 3000 rpm for 5 min between washes, and resuspended in wash buffer. Fluorescence was analyzed immediately by flow cytometry to detect active Caspase-3, with samples kept on ice.

#### Annexin V/7-AAD apoptosis assay by flow cytometry

To quantify apoptosis, 4T1-*CERS2*^+/+^ and 4T1-*CERS2*^−/−^ cells were treated with DH20931 at concentrations of 5, 10, and 20 µM for 24 h. Following treatment, apoptosis was assessed using the APC Annexin V Apoptosis Detection Kit with 7-AAD following the manufacturer’s protocol. Briefly, both floating and adherent cells were harvested, washed twice with ice-cold phosphate-buffered saline (PBS), and then resuspended in 1× Annexin V Binding Buffer at a concentration of 1 × 10⁶ cells/mL.

A 100 µL volume of the cell suspension (containing 1 × 10⁵ cells) was transferred to a 5 mL polystyrene tube. Five microliters (5 µL) of APC-conjugated Annexin V and 5 µL of 7-AAD viability dye were added to the cells. The samples were gently vortexed and incubated for 15 min at room temperature (25°C) in the dark. After incubation, 400 µL of 1× Binding Buffer was added to each tube to prepare the samples for immediate analysis.

Stained cells were analyzed on a BD FACSCanto™ II flow cytometer. The data were processed using FlowJo™ software to quantify the distribution of cells across different states. The cell populations were defined as follows: viable cells (Annexin V⁻/7-AAD⁻), early apoptotic cells (Annexin V⁺/7-AAD⁻) and late apoptotic/necrotic cells (Annexin V⁺/7-AAD⁺).

Quantitative data were statistically analyzed using GraphPad Prism (v9.0, GraphPad Software, San Diego, CA). The results were reported as the mean percentage of cells in each quadrant ± standard deviation (SD).

#### *In vivo* efficacy study

All animal procedures were performed in accordance with a protocol approved by the Institutional Animal Care and Use Committee (IACUC) of University of Florida (Protocol #202200000100). Once tumors were established and palpable, mice were randomly assigned to treatment groups (n = 8-10 mice per group). The treatment group received DH20931, administered at 6 mg/kg via intraperitoneal (*i.p.*) injection on a schedule of five times per week. DH20931 was dissolved in Captisol (1:1 mg/mg in 1 mL normal saline). Mice in the control group received the same amount of the Captisol. The study was concluded when tumors reached the predetermined endpoint of ≥ 15 mm in length. At this point, mice were euthanized by an approved method, and tumors and other relevant tissues were harvested for further analysis.

#### Molecular docking and binding free energy calculations

The three-dimensional structure of human Ceramide Synthase 2 was sourced from the AlphaFold Protein Structure Database (PDB ID: AF-Q3ZBF8-F1-model_v4). Due to the predictive nature of the model, the structure underwent a rigorous preparation protocol using the Protein Preparation Wizard in the Schrödinger Suite (release 2025.1, Maestro version 14.3; https://www.schrodinger.com) to ensure its suitability for subsequent molecular docking studies. The preparation process involved the following steps: assignment of bond orders, addition of hydrogen atoms, and generation of disulfide bonds. To address potential inaccuracies in the predicted model, missing side chains and loops were modeled and refined using Prime. The protonation states of ionizable residues at a physiological pH of 7.4 were predicted and optimized using PROPKA, followed by an optimization of the hydrogen-bonding network. All water molecules were removed, and the final structure was subjected to a restrained energy minimization using the OPLS3e force field until the root-mean-square deviation (RMSD) of heavy atoms converged to 0.30 Å. This minimization step serves to relieve any steric clashes and refine the geometry to a more energetically favorable conformation.^69^

The ligand was prepared for docking using the LigPrep module of the Schrödinger Suite. The OPLS3e force field was employed to generate low-energy 3D conformations of the ligand. This process included the exploration of possible ionization states at the target pH of 7.4 (± 2.0 pH units) using Epik, as well as the generation of relevant tautomers and stereoisomers to ensure a comprehensive representation of the ligand’s potential binding modes (Schrodinger Release 2023-24: LigPrep; Schrödinger, LLC: NewYork, NY, 2024; (https://www.schrodinger.com/platform/products/ligprep/#:∼:text=Overview,of%20the%20molecule%20in%203D).

To account for the flexibility of the binding site upon ligand binding, induced fit docking (IFD) was performed using the Glide and Prime modules of the Schrödinger Suite. The binding site was identified using SiteMap, and a docking grid was generated centered on the predicted active site. The IFD protocol was executed with default parameters, which involves an initial softened-potential docking of the ligand into a rigid receptor, followed by a refinement of the binding pocket residues within a defined cutoff of the ligand pose. The final ligand pose is then redocked with higher precision.^70^

Following the IFD-docking, the binding free energy of the protein-ligand complex was calculated using the Prime Molecular Mechanics/Generalized Born Surface Area (MM/GBSA) method. This approach computes the free energy of binding by combining the molecular mechanics energies with solvation energies calculated using the VSGB solvation model. The resulting ΔG(kcal/mol) provides an estimation of the binding affinity of the ligand for Ceramide Synthase 2.

#### Molecular dynamics simulation (MDS)

To investigate the dynamic stability of the docked protein-ligand complex, a 100-nanosecond (ns) molecular dynamics simulation was performed using the Desmond package in the Schrödinger Suite (Schrodinger Release 2024: Desmond Molecular DynamicsSystem; D. E. Shaw Research: New York, NY, 2024; https://www.schrodinger.com/life-science/download/release-notes/release-2024-1/).

The docked complex from the IFD calculations served as the initial coordinates for the MD simulation. The system was prepared using the System Builder tool. The complex was solvated in an orthorhombic box with SPC (Simple Point Charge) water molecules, extending at least 10 Å from the protein surface in all directions to ensure no direct interactions with its periodic image. The system was neutralized by the addition of Na^+^ counterions. The OPLS3e force field was used for all atoms in the system.^71^

The prepared system underwent a default equilibration protocol within Desmond. This multi-stage process involves a series of restrained minimizations and short MD simulations under NVT (constant volume and temperature) and NPT (constant pressure and temperature) ensembles to gradually relax the system and bring it to the desired temperature (300 K) and pressure (1 bar). Following equilibration, the production MD simulation was carried out for 100 ns in the NPT ensemble, with temperature and pressure maintained using the Nosé-Hoover chain thermostat and Martyna-Tobias-Klein barostat, respectively.

The stability of the protein-ligand complex throughout the simulation was assessed by analyzing the trajectory. The root-mean-square deviation (RMSD) of the protein backbone and the ligand heavy atoms were calculated to evaluate conformational changes. The root-mean-square fluctuation (RMSF) of individual residues was computed to identify flexible regions of the protein. The Simulation Interaction Diagram tool was used to analyze the evolution of protein-ligand interactions, such as hydrogen bonds, hydrophobic contacts, and water bridges, over the course of the simulation.

#### *In silico* ADME analysis

The ADME properties of DH20931 were evaluated using the Swiss-ADME online software (http://www.swissadme.ch/index.php). The results, summarized in Table 1, indicate that the molecule adheres to key drug-likeness criteria. The molecular weight of the DH compound falls within the acceptable range, and its topological polar surface area (TPSA) was calculated to be 27.27 Å², which supports good membrane permeability. Additionally, the compound’s LogS and LogP values suggest favorable aqueous solubility and lipophilicity, respectively. Importantly, the molecule showed no violations of Lipinski’s Rule of Five and did not trigger any PAINS (Pan-Assay Interference Compounds) alerts, further supporting its potential as a drug-like candidate.

#### *In vitro* ADME analysis

##### Pharmacokinetic study

A pharmacokinetic study of DH20931 was performed after intraperitoneal administration (10 mg/kg) in male mice (25 g of body weight). DH20931 was dissolved in Captisol (1:1 mg/mg in 1 mL normal saline). During the pharmacokinetic study, animals were allowed free access to food and water. Blood (80 μL) was collected in heparinized tubes at pre-dose, and 0.25, 0.5, 1, 2, 4, and 8 h, post-dose.

##### Bioanalysis

Plasma and tissue homogenate samples (20 μL) underwent protein precipitation using 80 μL of acetonitrile containing an internal standard (IS). After quenching, samples were vortex mixed, filtered, centrifuged, and injected onto a Waters Acquity UPLC system coupled with a Xevo® TQ-S micro triple quadrupole mass spectrometer. Source and compound parameters for DH20931 estimation are detailed in Table S5. Chromatographic separation was achieved on a Waters Acquity UPLC BEH C18 column (1.7 µm, 2.1 × 100 mm) with a VanGuard pre-column, using a linear gradient: 0–0.3 min, 20% B; 0.3–1.3 min, 20–85% B; 1.3–1.5 min, 85% B; 1.5–2.0 min, 85-20% B.

Mobile phase A consisted of 0.1% formic acid in water, and mobile phase B was acetonitrile. Calibration and quality control standards were prepared by spiking known concentrations of DH20931 into blank plasma. The concentration-time data were analyzed using non-compartmental analysis with Phoenix® software (Certara USA, Inc., Princeton, NJ).^72^

##### Caco-2 permeability

Caco-2 cells (passage 27) were cultured in T-75 flasks with DMEM supplemented with 10% FBS, 1% nonessential amino acids, 2 mM L-glutamine, 100 μg/mL streptomycin, and 100 U/mL penicillin, and maintained at 37°C with 5% CO₂. Cells were seeded onto 24-well Transwell® inserts (0.33 cm², 0.4 μm pores) at a density of 0.27×106 cells/cm². Media was changed every other day for 14 days, then daily until day 23. Before the assay, DMEM was replaced with pH 7.4 HBSS supplemented with dextrose and HEPES. Monolayer integrity was confirmed by Transepithelial/endothelial electrical resistance (TEER) values greater than 400 Ω·cm². Atenolol and propranolol served as low and high permeability markers, respectively. Digoxin and sulfasalazine were used as P-glycoprotein (P-gp) and breast cancer resistance protein (BCRP) substrates, respectively. Bidirectional transport studies (A-to-B and B-to-A) were performed using 10 μM solutions of test compounds and controls. Experiments were also conducted in the presence of P-gp and BCRP inhibitors to evaluate changes in the efflux ratio and determine their roles in B-to-A efflux. Samples were collected from the acceptor chamber at 0 and 2 h, quenched with acetonitrile containing IS (5 ng/mL), and filtered using a solvinert 96-well filter plate with 0.45 µm pore size (Millipore, St. Louis, Missouri) by centrifuging at 2000 rpm for 5 min at 4°C. Samples were then analyzed by ultra-performance liquid chromatography−tandem mass spectrometry (UPLC-MS/MS).^72^

##### Metabolic stability

Metabolic stability was assessed using rat liver microsomes. The test compound DH209431 (1 μM) was incubated with 1 mg/mL microsomal protein in 50 mM phosphate buffer (pH 7.4), with verapamil as a positive control. The reaction was initiated by adding 1 mM NADPH, a required cofactor for cytochrome P450 activity. Incubations were carried out at 37°C in a 5% CO₂ atmosphere at 120 rpm. Aliquots were collected at 0, 5, 10, 15, 30, 45, and 60 min. Reactions were stopped by adding ice-cold acetonitrile containing IS. Samples were processed via protein precipitation, filtered, and analyzed as described in the bioanalysis section. Area ratios (compound/internal standard) were used to calculate the percentage of compound remaining at each time point. The elimination rate constant (k) and half-life (T_1/2_) were derived from the slope of the natural log-transformed area ratio versus time plot.^73^

##### Plasma-protein binding

The fraction of unbound DH20931 in human plasma was determined using an equilibrium dialysis assay.^72^ This technique is crucial for assessing pharmacologically active drug levels, as only the unbound fraction contributes to efficacy and toxicity. A 96-well Teflon dialysis plate (HTDialysis, Gales Ferry, CT, USA) with 12–14 kDa molecular weight cutoff cellulose membranes was used. Membranes were preconditioned sequentially in LC-MS grade water (60 min), 20% ethanol (20 min), and 50 mM phosphate buffer (pH 7.4). Plasma samples were preincubated at 37°C, 5% CO₂ for 15 minutes, then spiked with 1 μM DH20931 and control compounds (ketoconazole and atenolol). Initial aliquots were taken for baseline measurements. For dialysis, 120 μL of spiked plasma was added to the donor side, and 120 μL of phosphate buffer was added to the receiver side. Plates were sealed and incubated at 37°C, 120 rpm for 6 h. After dialysis, 20 μL from each side was matrix-matched and quenched with acetonitrile containing IS. Samples were filtered and analyzed as described previously. Area ratios were used to calculate percent unbound and percent recovery, following the protocol reported by Gour et al.^72^

## QUANTIFICATION AND STATISTICAL ANALYSIS

Data are presented as the mean ± standard deviation (SD) from at least three independent experiments unless otherwise stated. Statistical analyses were performed using GraphPad Prism 9 or SigmaPlot 15. Specific quantification methods are as follows:

### Cell viability

IC_50_ values were determined by fitting dose-response curves using non-linear regression.

### Clonogenic and Spheroid assays

Colonies and spheroid areas were quantified using ImageJ software (NIH).

### Western blots

Densitometry of protein bands was performed using ImageJ.

### Fluorescence and microscopy

Fluorescence intensity from activity assays, Ca²⁺ imaging, and immunofluorescence was quantified using ImageJ or native microscope software (e.g., ImageQuant, NIS Elements). ER-mitochondria co-localization was quantified using the JACoP plugin in ImageJ to calculate Mander’s overlap coefficients.

### Flow cytometry

Data were analyzed using FlowJo software to quantify percentages of apoptotic cells.

### Statistical significance

Comparisons between two groups were performed using two-tailed, unpaired Student’s t-tests. Comparisons among multiple groups were performed using one-way analysis of variance (ANOVA) followed by an appropriate post-hoc test. A p-value of < 0.05 was considered statistically significant.

## SUPPLEMENTARY FIGURES AND TABLES

**Figure S1.**
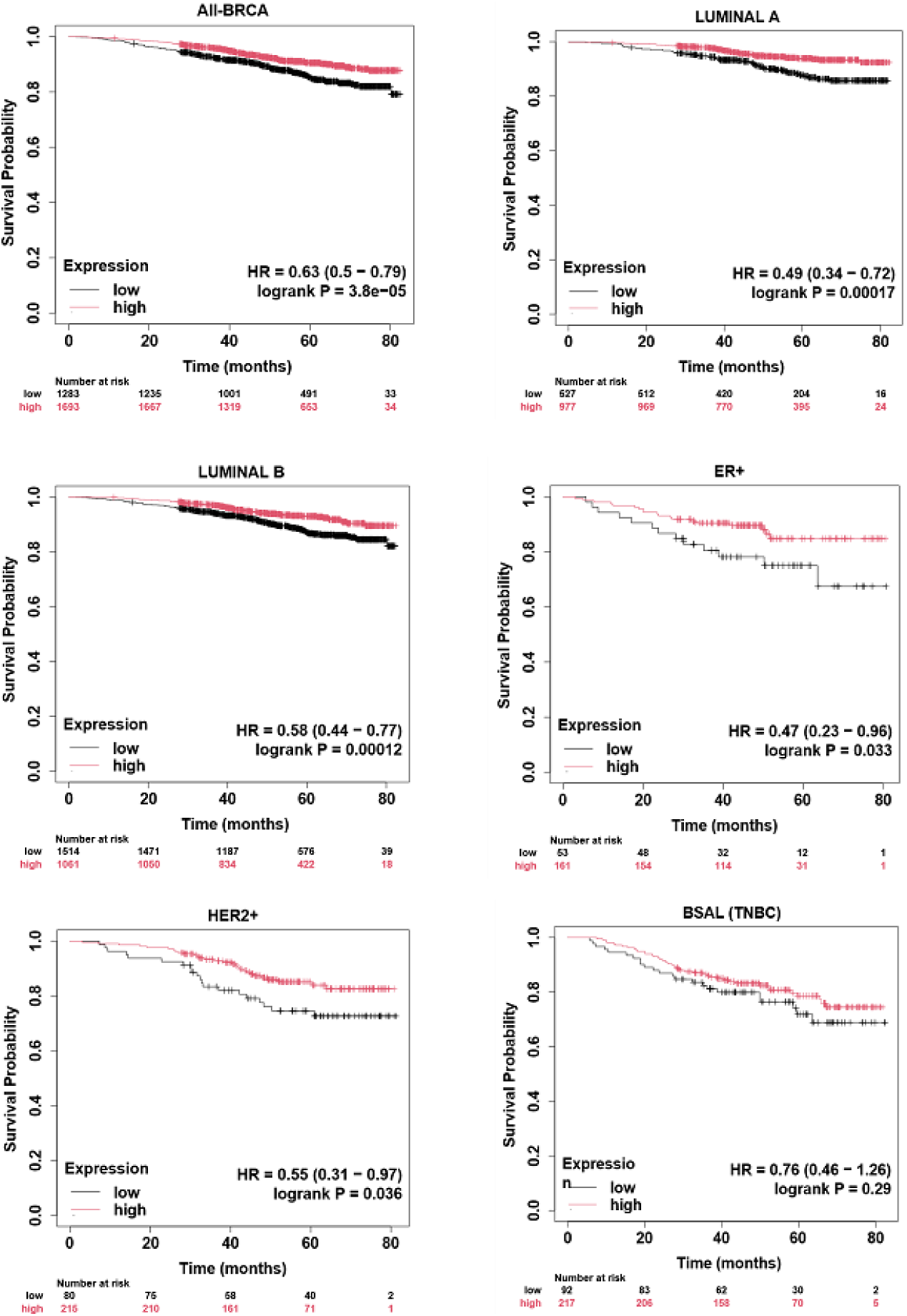
*CERS2* expression is correlated with longer survival of breast cancer patients. Kaplan-Meier Plots based on breast cancer RNA=seq data from All-BRCA, Luminal A, Luminal B, ER+, HER2+ and Basal (TNBC) tumor samples. HR: hazard ratio, BRCA: breast cancer.

**Figure S2.**
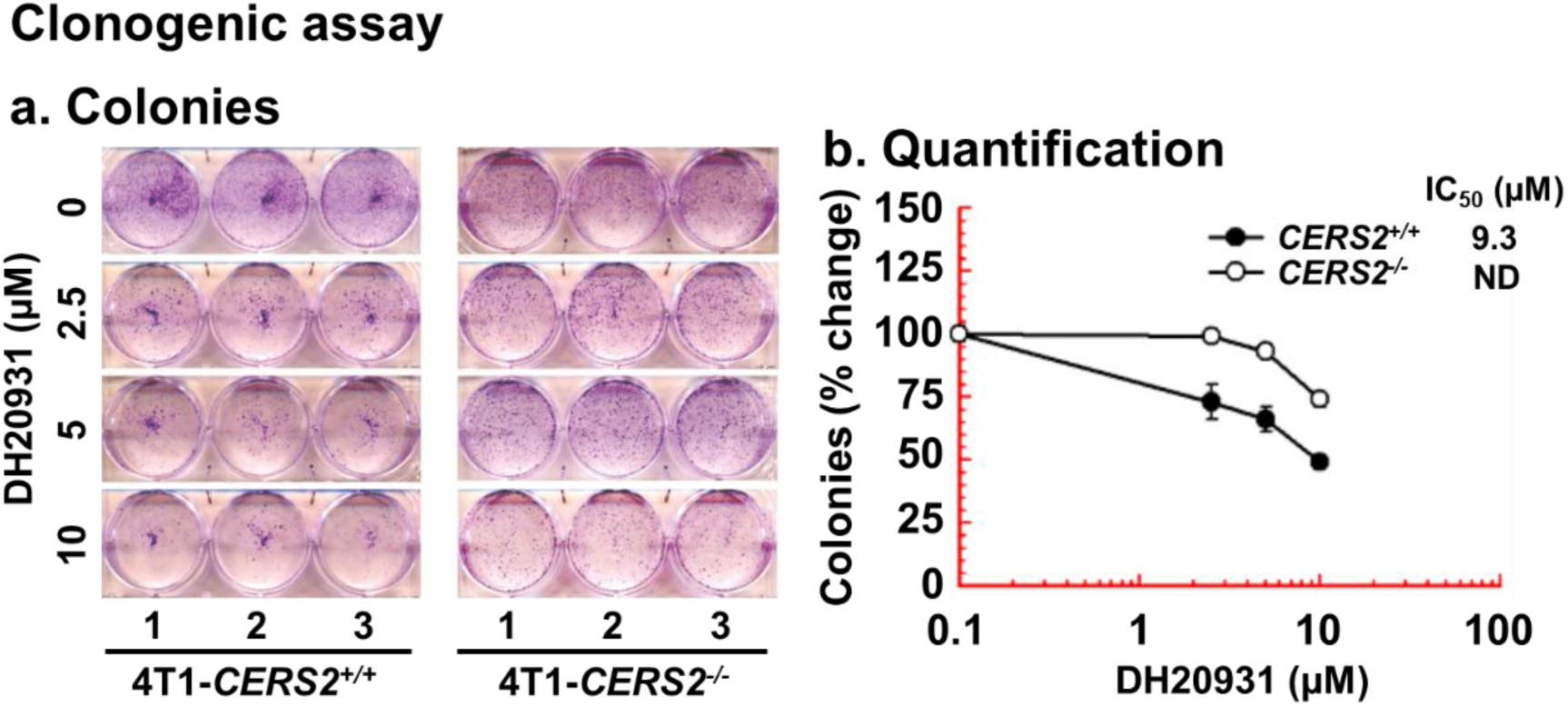
DH20931 reduces the clonogenic activity of 4T1 cells in a *CerS2*-dependent manner. 4T1-*CERS2*⁺^/^⁺ and 4T1-*CERS2*⁻^/^⁻ cells were treated with the indicated concentrations of DH20931 for 7 days. **(A)** Representative images of colonies stained with crystal violet. **(B)** Colony numbers were quantified using ImageJ software and plotted. Data are presented as the mean ± SEM from three independent replicates. ND = not detected.

**Figure S3.**
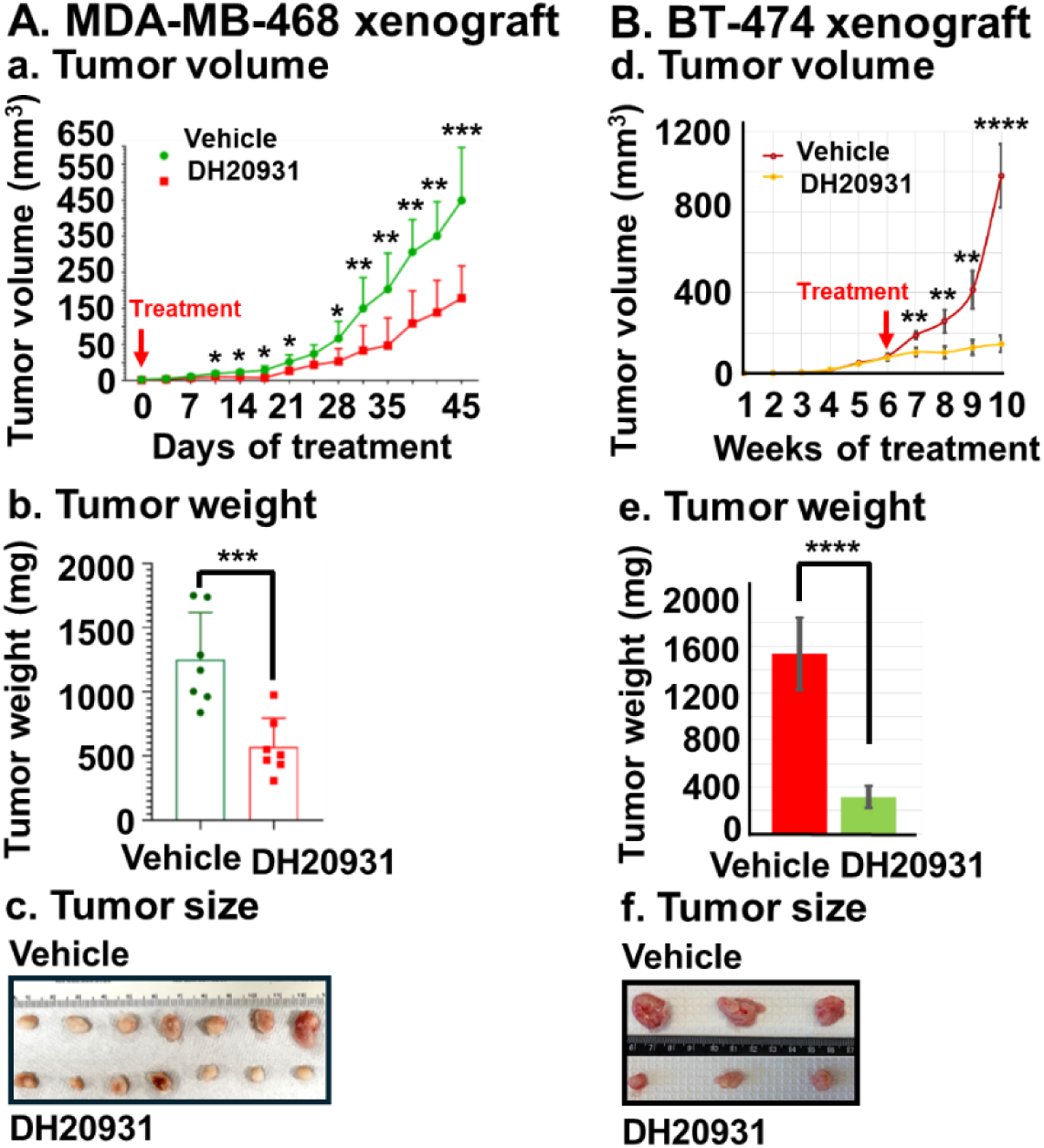
DH20931 demonstrates antitumor activity in orthotopic xenograft models of TNBC (MDA-MB-468) and Luminal B (BT-474) breast cancer. **(A)** DH20931 suppresses MDA-MB-468 (TNBC) tumor growth in an orthotopic model. NSG (NOD.Cg-*Prkdc*^scid^ *Il2rg*^tm1Wjl^/SzJ) female mice (n=7/group) were treated with vehicle or DH20931 (6 mg/kg, 5 days/week). Graphs depict **(a)** tumor volume over time, **(b)** final tumor weight, and **(c)** representative images of excised tumors. Data are presented as mean ± SD. *p* < 0.05, \**p* < 0.01, \*\**p* < 0.001. **(B)** DH20931 suppresses BT474 (Luminal A) tumor growth in an orthotopic model. Mice (n=3/group) were treated with vehicle or DH20931 (6 mg/kg, 5 days/week). Graphs depict **(a)** tumor volume over time, **(b)** final tumor weight, and **(c)** representative images of excised tumors. Data are presented as mean ± SD. \**p* < 0.01, \*\*\**p* < 0.0001.

**Figure S4.**
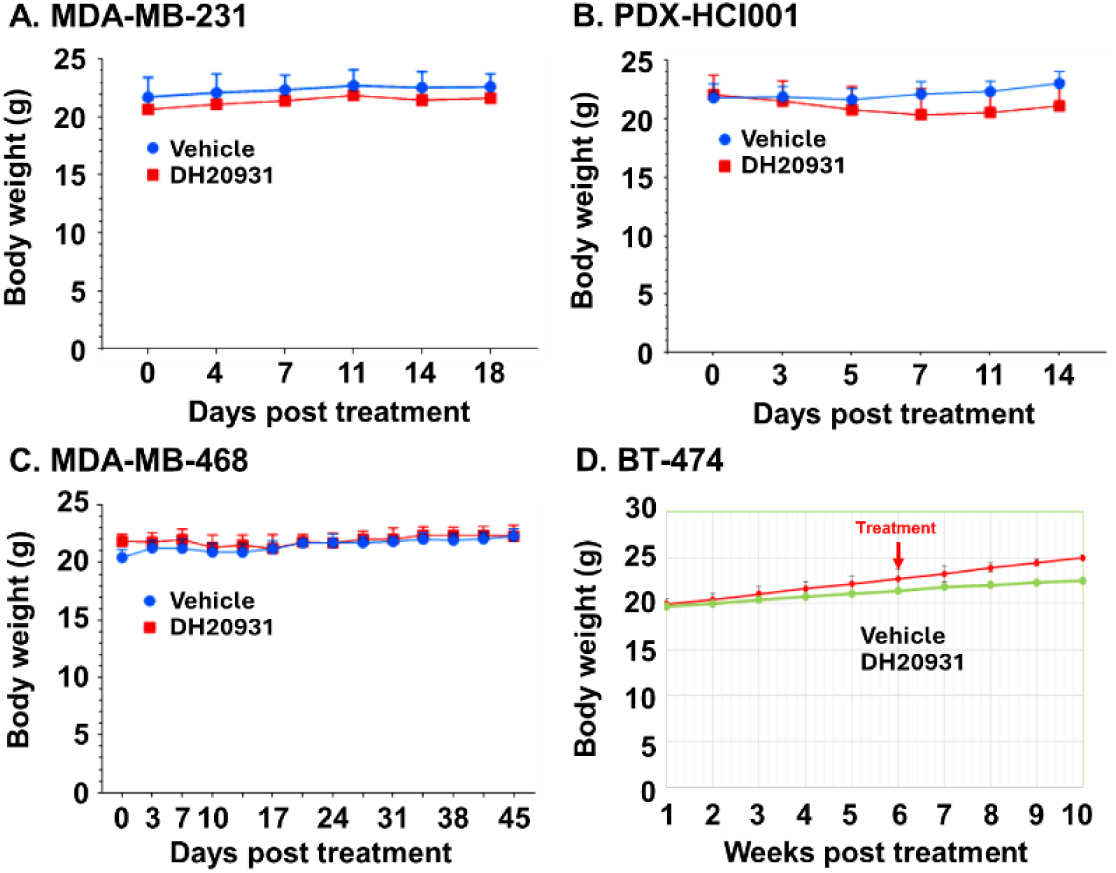
Body weight of female NSG female mice bearing MDA-MB-231, PDX-HCI001, MDA-MB-468 and BT-474 tumors. Data are presented as Mean ± SD of 3-11 mice in each group, as shown in Figure 3 and Figure S3.

**Figure S5.**
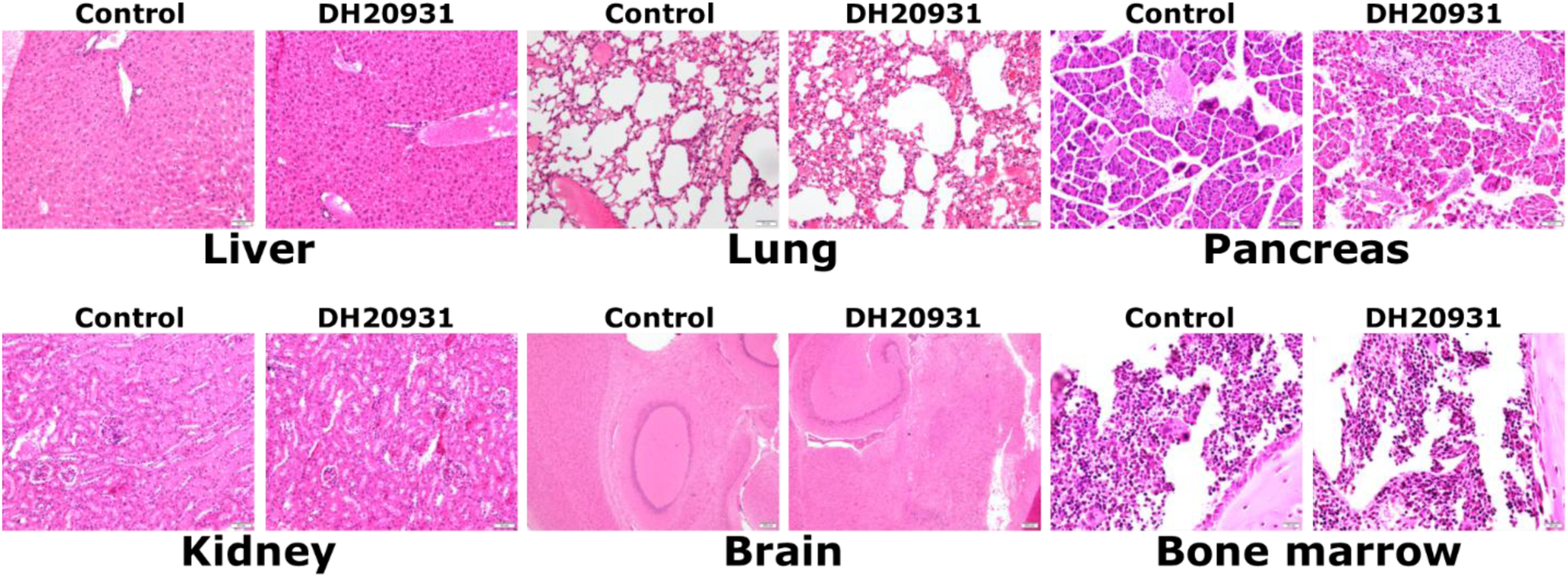
DH20931 treatment does not exhibit *in vivo* toxicity. Mice were treated intraperitoneally (*i.p.*) with vehicle or DH20931 (6 mg/kg) daily for 21 days. On day 21, mice were euthanized, and major organs were collected for histological examination. Throughout the study, all mice remained healthy and exhibited normal characteristics (e.g., body weight gain, movement, fur). Histological analysis of H&E-stained organ sections revealed no significant pathological differences between control and DH20931-treated mice, indicating a favorable safety profile.

---

**Table S1.** Mean concentrations of different blood cell count in female NSG mice in control and DH20931 treatment for 21 days, five days per week (Monday-Friday).

| **Blood Cells** | **Control** | **DH20931**<br>**(7 day)** | **DH20931**<br>**(14 day)** | **DH20931**<br>**(20 day)** |
| --- | --- | --- | --- | --- |
| White Blood Cells (10<sup>3</sup>/uL) | 1.89 ± 0.28 | 3.36 ± 1.57 | 2.13 ± 0.25 | 2.52 ± 0.87 |
| Neutrophils (10<sup>3</sup>/uL) | 1.22 ± 0.22 | 2.08 ± 1.22 | 1.46 ± 0.58 | 1.67 ± 0.47 |
| Lymphocytes (10<sup>3</sup>/uL) | 0.16 ± 0.17 | 0.36 ± 0.22 | 0.17 ± 0.24 | 0.24 ± 0.33 |
| Monocytes (10<sup>3</sup>/uL) | 0.29 ± 0.03 | 0.42 ± 0.19 | 0.30 ± 0.09 | 0.35 ± 0.08 |
| Eosinophils (10<sup>3</sup>/uL) | 0.166 ± 0.14 | 0.41 ± 0.38 | 0.14 ± 0.13 | 0.20 ± 0.18 |
| Basophils (10<sup>3</sup>/uL) | 0 ± 0 | 0 ± 0 | 0 ± 0 | 0 ± 0 |
| Neutrophils (%) | 65.54 ± 11.85 | 62.20 ± 18.39 | 67.70 ± 25.37 | 67.56 ± 10.58 |
| Lymphocytes (%) | 8.24 ± 7.46 | 11.12 ± 6.98 | 8.72 ± 12.40 | 6.18 ± 7.97 |
| Monocytes (%) | 15.54 ± 3.45 | 12.50 ± 0.84 | 14.30 ± 3.23 | 14.94 ± 4.98 |
| Eosinophils (%) | 8.44 ± 5.82 | 11.60 ± 9.17 | 7.16 ± 6.64 | 7.32 ± 5.77 |
| Basophils (%) | 2.24 ± 3.16 | 2.57 ± 3.02 | 0 ± 0 | 4.00 ± 2.69 |
| Red Blood Cells (10<sup>6</sup>/uL) | 8.52 ± 0.30 | 8.60 ± 0.47 | 8.31 ± 0.55 | 8.84 ± 0.39 |
| Hemoglobin (g/dL) | 14.10 ± 0.74 | 13.60 ± 1.00 | 13.38 ± 0.59 | 13.92 ± 0.16 |
| Hematocrit (%) | 41.56 ± 1.75 | 41.32 ± 2.33 | 39.58 ± 3.26 | 43.14 ± 0.85 |
| Mean Corpuscular Volume (fL) | 48.76 ± 0.51 | 48.05 ± 0.20* | 47.58 ± 0.87* | 48.86 ± 1.86 |
| Mean Corpuscular Hemoglobin (pg) | 16.56 ± 0.41 | 15.8 ± 0.42* | 16.12 ± 0.43 | 15.74 ± 0.85 |
| Mean Corpuscular Hemoglobin Concentration (g/dL) | 33.96 ± 0.66 | 32.87 ± 0.88 | 33.92 ± 1.52 | 32.22 ± 0.74** |
| Red Cell Distribution Width – Coefficient of Variation (%) | 14.76 ± 0.47 | 15.22 ± 0.57 | 17.56 ± 4.10 | 15.00 ± 0.36 |
| Platelets(10<sup>3</sup>/uL) | 1215 ± 183 | 1327 ± 278 | 1171 ± 229 | 1279 ± 239 |
| Mean Platelet Volume (fL) | 5.4 ± 0.12 | 5.37 ± 0.12 | 5.24 ± 0.48 | 5.58 ± 0.17 |

Female NSG mice were treated with DH20931 (6 mg/kg body weight in 1:1 Captisol/normal saline) for 5 days/week (Monday-Friday). Blood samples were collected at 0, 7, 14 and 20 days in heparinized tubes and immediately processed for complete blood cell counts (CBC) using a HESKA Element HT5 Veterinary Hematology Analyzer. Data are presented as mean ± SEM of 4-5 samples in each group. \**p* < 0.05, \*\**p* < 0.01.

**Table S2.** *In silico* ADME and physicochemical properties of DH20931.

| **Name** | **Mol. Wt**<br>**(350-500**<br>**g/mol)** | **TPSA**<br>**(7-200 Å)** | **LogS**<br>**(-6.5-0.5)** | **Lipinski**<br>**Violation** | **PAINS** | **logP**<br>**(2-5)** |
| --- | --- | --- | --- | --- | --- | --- |
| DH20931 | 333.40 | 27.27 | -3.60 | 0 | 0 | 2.52 |

The table presents key physicochemical features relevant to drug-likeness, including molecular weight, topological polar surface area (TPSA), predicted aqueous solubility (LogS), and lipophilicity (LogP). The analysis also includes compliance with Lipinski's Rule of Five and flags for potential Pan-Assay Interference Compounds (PAINS).

**Table S3.** PAMPA permeability studies of marker and DH20931.

| Markers and test compounds | Permeability ($P_{app}$) ($\times 10^{-6}$ cm/sec) | Permeability class (High/low) |
| --- | --- | --- |
| DH20931 | $0.007 \pm 0.00$ | Low |
| Propranolol | $11.15 \pm 2.87$ | High |
| Caffeine | $11.93 \pm 1.41$ | High |
| Atenolol | $0.008 \pm 0.005$ | Low |

Data are presented as Mean ± SD of three estimations.

**Table S4.**
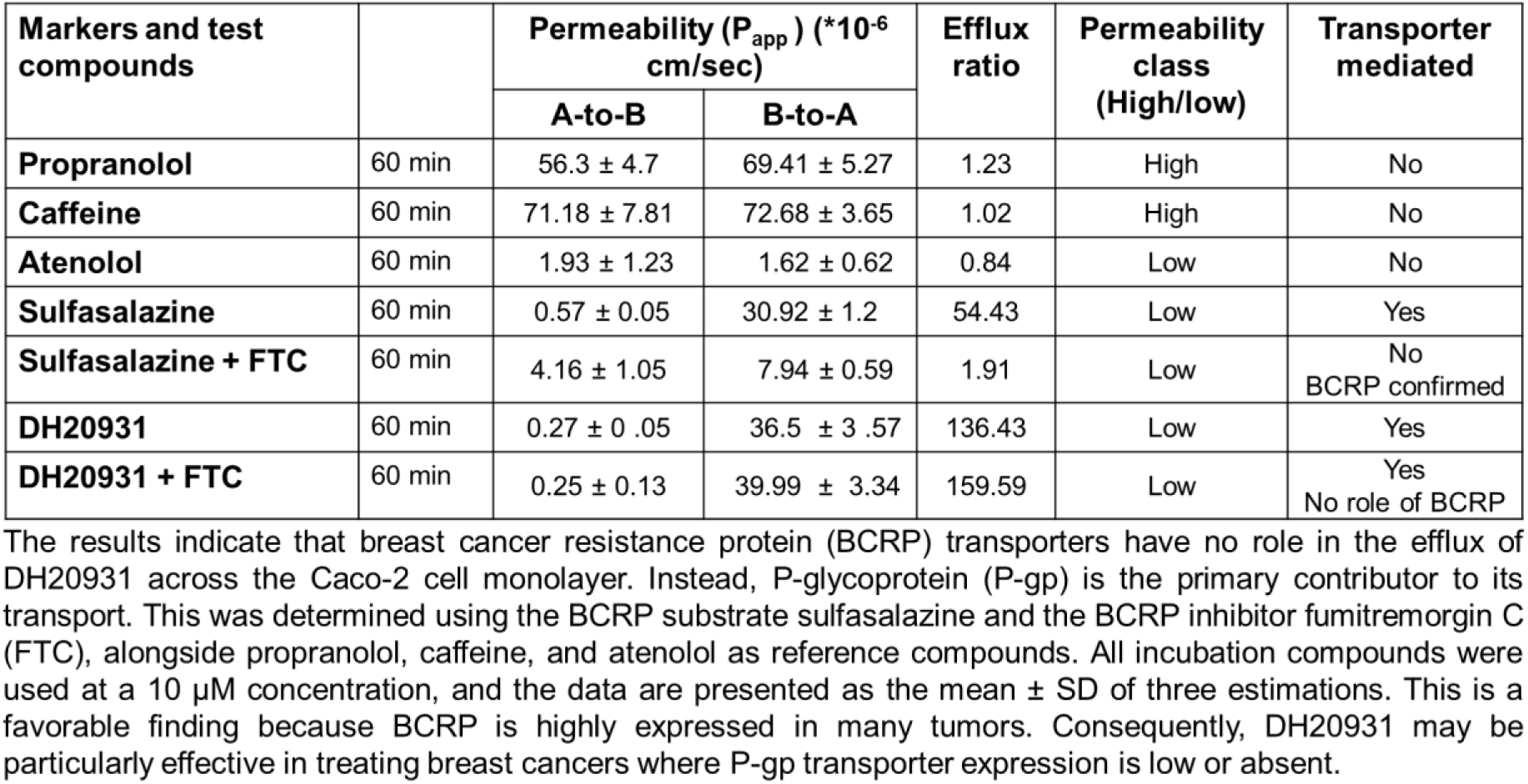
Caco-2 permeability studies of marker and test compounds.

**Figure S6.**
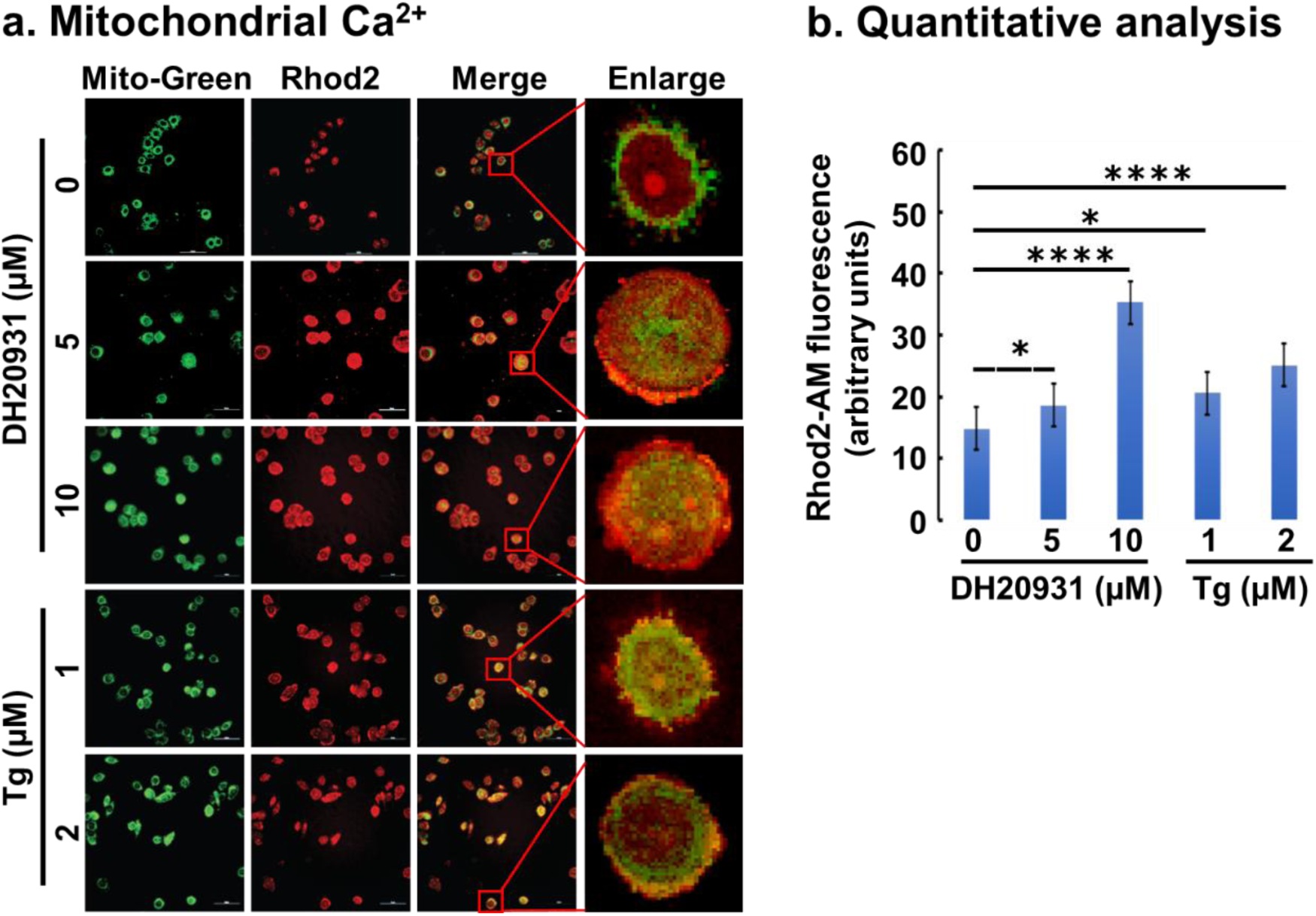
DH20931 increases mitochondrial Ca²⁺ levels in MDA-MB-468 cells. MDA-MB-468 cells were treated with DH20931 for 6 h and co-loaded with Mag-Fluo-4 (ER Ca²⁺ indicator) and Rhod-2/AM (mitochondrial Ca²⁺ indicator). **(A)** Representative confocal microscopy images visualizing intracellular Ca²⁺ localization. **(B)** Quantification of mitochondrial Ca²⁺ fluorescence intensity. Data were measured using ImageJ and are presented as the mean ± SEM from 30 cells per group. \*\*\**p* < 0.0001.

**Figure S7.**
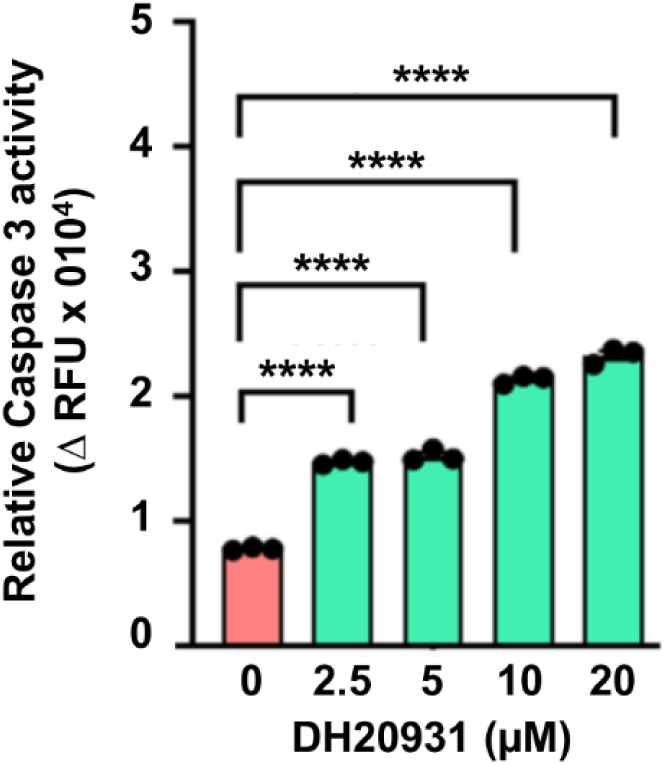
DH20931 treatment induces Caspase-3 activity in MDA-MB-468 cells. Cells treated with the indicated concentrations of DH20931 for 24 h. Activity was measured using a Caspase-3 activity assay kit. Data are presented as mean ± SEM of three independent experiments. ****p < 0.0001.

**Table S5.** Source and compound parameters for estimation of DH20931 using UPLC-MS/MS.

| Parameters | Value |
| --- | --- |
| MS/MS Conditions |  |
| Scan type | Multiple reaction monitoring (MRM) |
| Source | Electrospray ionization (ESI) |
| Ion polarity | Positive |
| Capillary voltage (kV) | 0.5 |
| Source temperature (°C) | 150 |
| Desolvation temperature (°C) | 450 |
| Desolvation gas (L/h) | 900 |
| Cone gas flow rates (L/h) | 50 |
| Cone voltage for DH20931 (V) | 34 |
| Cone voltage for internal standard (IS) (V) | 34 |
| Collision energy for DH20931 (V) | 66 |
| Collision energy IS (V) | 20 |
| Ion transition for DH20931 (*m/z*) | 333.2 > 245.2 |
| Ion transition for phenacetin (IS, *m/z*) | 180.1 > 110.1 |

